# Streamlined and sensitive mono- and di-ribosome profiling in yeast and human cells

**DOI:** 10.1101/2023.02.01.526718

**Authors:** Lucas Ferguson, Heather E. Upton, Sydney C. Pimentel, Amanda Mok, Liana F. Lareau, Kathleen Collins, Nicholas T. Ingolia

## Abstract

Ribosome profiling has unveiled diverse regulations and perturbations of translation through a transcriptome-wide survey of ribosome occupancy, read out by sequencing of ribosome-protected mRNA fragments. Generation of ribosome footprints and their conversion into sequencing libraries is technically demanding and sensitive to biases that distort the representation of physiological ribosome occupancy. We address these challenges by producing ribosome footprints with P1 nuclease rather than RNase I and replacing RNA ligation with Ordered Two-Template Relay, a single-tube protocol for sequencing library preparation that incorporates adapters by reverse transcription. Our streamlined approach reduced sequence bias and enhanced enrichment of ribosome footprints relative to ribosomal RNA. Furthermore, P1 nuclease preserved a myriad of distinct juxtaposed ribosome complexes informative about yeast and human ribosome fates during translation initiation, stalling, and termination. Our optimized methods for mRNA footprint generation and capture provides a richer translatome profile using lower input and fewer technical challenges.

## Introduction

Ribosome profiling provides a global, high-resolution view of translation by deep sequencing of ribosome-protected mRNA fragments (RPFs). By counting the number of RPFs along each protein coding sequence (CDS), the rate of protein synthesis can be estimated across the proteome at the point of cell lysis. Careful analysis of the position of each RPF enables deeper interpretations of ribosome occupancy, such as the decoding rate of individual codons. This approach has yielded insights about translation mechanisms, non-canonical events such as frameshifting and reinitiation, and open reading frame (ORF) discovery [1, 2, 3, 4, 5, 6, 7, 8]. Further insights have been gleaned from adaptations of ribosome profiling that capture specific ribosome sub-populations or pre-initiation complexes [9, 10, 11, 12, 13, 14, 15]. Despite these benefits, widespread use of ribosome profiling has been hampered by labor-intensive workflows with high RNA input requirements and low information return due to excessive capture of ribosomal RNA (rRNA) fragments generated during obligatory nuclease digestion [1].

Ribosome profiling has two essential steps: treating cell lysate with a nuclease that degrades accessible mRNA to generate RPFs, and converting these RPFs into a complementary DNA (cDNA) library for high-throughput sequencing. Each step poses distinctive technical challenges. The ideal nuclease would cleave unprotected mRNA with no sequence bias while sparing the structural RNA of the ribosome. The original and most widely used nuclease, *Escherichia coli* RNase I, has no nucleotide preference but readily digests rRNA, leading to substantial rRNA contamination of RPFs [1].

Alternatives such as micrococcal nuclease (MNase), RNase A, and RNase T1 can reduce rRNA contamination, but they have strong RNA sequence preferences so they do not provide the single-nucleotide resolution of RNase I RPFs and may distort measurements of translation speed and ribosome stalling [16]. Likewise, the ideal library generation approach would convert RPFs into a sequencing library efficiently and with little bias. Existing workflows join the 3′ ends of RPFs to adaptor oligonucleotides utilizing ligases, which are inefficient and have sequence biases [17]. Moreover, use of ligase obliges denaturing gel purifications to separate product from unreacted substrates, while also providing opportunities for nuclease contamination or stochastic RNA decay during elution at ambient temperatures.

We recently reported a technically straightforward and low-bias approach to construct small RNA sequencing libraries that takes advantage of the enzymatic activities of reverse transcriptase (RT) enzymes from eukaryotic retroelements. We used the terminal transferase and template-jumping cDNA synthesis activities of an engineered retroelement RT to develop an Ordered Two-Template Relay (OTTR) protocol for small RNA library generation [18]. This rapid and single-tube workflow eliminates any need for RNA or DNA ligases, replacing them with a programmed sequence of template jumps that directs continuous cDNA synthesis from a 5′ adaptor duplex primer across the input RNA template and then a 3′ adaptor template [18].

Here we describe a comprehensive redevelopment of the widely used ligation-based ribosome profiling workflow which improves labor costs, input requirements, and time necessary to generate sequenceable libraries. We first established OTTR as a suitable replacement for RPF sequencing library generation by direct comparison with the ligation-based approach and an in-depth analysis of library generation artifacts [19]. We then developed and validated P1 nuclease — a sequence-independent, single-strand nuclease — as an alternative to RNase I for RPF production in yeast and human cell lysates. We found that P1 nuclease preserved ribosome integrity better than RNase I due to reduced rRNA cleavage, an improvement which reduced rRNA contamination in RPF libraries. P1 nuclease footprinting and OTTR library synthesis detected the ribosomal stalling induced by impaired translation, and further revealed several configurations of adjacent, collided ribosomes (disomes), which indicate stalls in protein synthesis and trigger ribosome and/or mRNA turnover by ribosome quality control (RQC) pathways. Lastly, we further streamlined P1+OTTR ribosome profiling by eliminating any requirement for gel electrophoresis in RPF recovery, instead using a small RNA enrichment approach, mirRICH. Taken together this dramatically simplified workflow will aid in the broader adoption of ribosome profiling.

## Results

### OTTR is a favorable alternative to ligation-based workflows in ribosome profiling

To assess whether OTTR library generation is beneficial for ribosome profiling, we compared it to a standard ligation-based workflow [19]. We prepared a single pool of *Saccharomyces cerevisiae* (budding yeast) RPFs by RNase I digestion and split this sample into two portions to compare the two workflows. We generated OTTR ribosome profiling libraries from these footprints in a single-tube OTTR reaction over the course of about 4 hours, and we created ligation-based libraries for comparison by following the typical 3-day ligation-based workflow (**Figure 1a**). Because OTTR is capable of generating libraries from very low input, we used tenfold less RPFs for OTTR library generation than for the ligation-based workflow. The ligation-based method converted ∼400 ng of input RNA to 24 nmoles of DNA library while OTTR converted ∼40 ng to 180 nmoles of library, a 75-fold improvement. Further, the OTTR library contained a lower fraction of unwanted library generation byproducts such as adapter-only reads (**Extended Data 1a**). The RPF-derived reads in the two libraries were highly correlated at both gene-occupancy and codon-occupancy levels (**Figures 1b-c**). Thus, OTTR provides comparable data but with improved RPF conversion efficiency and higher yield of biologically interpretable reads than the ligation-based workflow.

**Figure 1:**
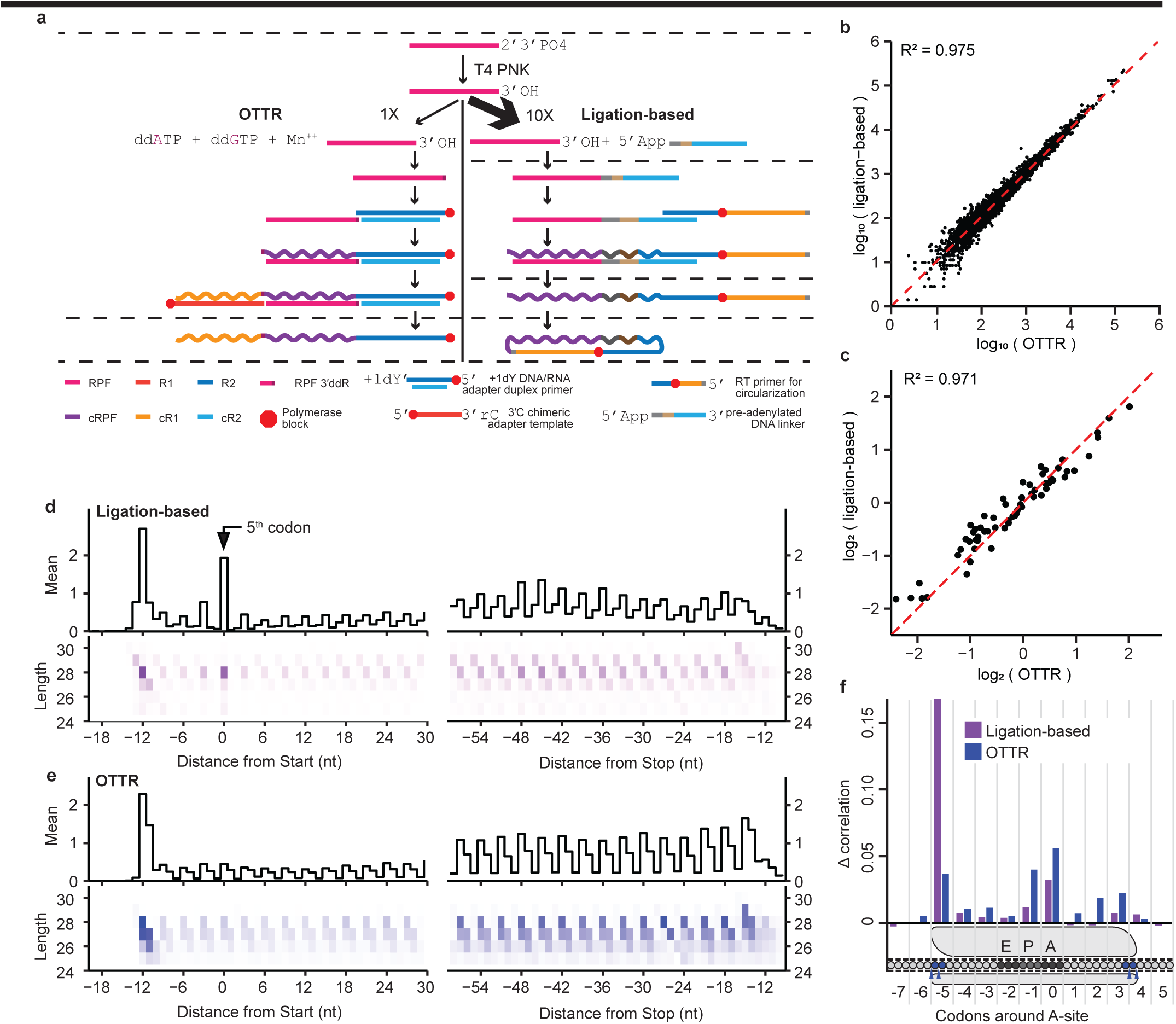
Highly correlated ribosome profiles derived from ligation-based or OTTR library generation. **a**, Schematic of library generation by OTTR or ligation-based protocols from a single pool of RNase I derived RPFs (pink), split unevenly. Dashed lines between steps indicate a size-selection step by denaturing urea polyacrylamide gel electrophoresis (PAGE), nucleic acid elution, and precipitation. In OTTR, the Illumina read 2 primer sequence (R2, DNA) with a single 3′dT or 3′dC overhang is annealed to a partial complementary R2 sequence (cR2, RNA). This +1 3′dY DNA/RNA adapter primer duplex captures a 3′ddR labeled RPF (RPF 3′ddR) and primes synthesis of cDNA (cRPF). Non-templated dG addition to the cRPF strand enables capture of the 3′rC chimeric adapter template, which encodes the Illumina read 1 primer sequence (R1 RNA-DNA chimera). Following cRF extension by synthesis of the adapter-template complement strand (cR1), subsequent template jumps are prevented by a 5′Cy5 modification on the adapter template (red octagon). **b**, Comparison of gene-level ribosome occupancy estimates from libraries generated in (a). Read counts are for RPFs aligned to verified CDSs excluding those RPFs that are aligned to the first 15 and last 10 codons. **c,** Comparison of mean codon-level occupancy estimates from libraries generated in (a). Aligned RPFs were assigned to an A-site codon and counted. These counts were then rescaled by the mean codon count for the gene, excluding those RPFs that are aligned to the first 15 and last 10 codons, and averaged across the translatome. **d-e,** Metagene averages around the start (left) and stop (right) codons for either (**d**) OTTR or (**e**) ligation-based libraries. Aligned RPFs for each CDS were first rescaled by the mean codon count for the gene, excluding those RPFs that are aligned to the first 15 and last 10 codons, and then averaged across the translatome. Footprints were tabulated according to either the 5′ aligned position alone (shown at top as a black line), or both 5′ aligned position and read length (shown at bottom as a matrix of distinct RPF lengths and positions). **f**, Per-codon contributions to iχnos machine learning models of RPF occupancy profiles. A model based on a widow of 13 codons (-7 to +5) around the A-site was compared with thirteen additional models, each omitting one codon from the model. The contribution of a codon position to RPF occupancy profile was inferred from the change in Pearson’s correlation coefficient between the predicted ribosome occupancy versus actual ribosome occupancy changed when the codon was omitted (Y-axis).

A strength of ribosome profiling is the ability to infer the codon being decoded by each ribosome based on the precise size and position of each RPF. The peptidyl tRNA (P) and acceptor tRNA (A) sites are typically offset 12 – 14 and 15 – 17 nucleotides (nt), respectively, from the 5′ end of an RNase I RPF. Coupled with the 3-nt periodicity of ribosome elongation, we can rely on the exact length and reading frame of an aligned RPF to determine the A-site position of each individual RFP [1]. In theory, these A-site offsets can be calibrated empirically from ribosome occupancy at initiation and termination codons (**Figure 1d**), but in practice, library generation artifacts must be considered when relating both length and reading frame of an RPF to its A-site offset. Here, we noted that RPF sequences from the OTTR library were slightly shorter than those from the ligation-based library, despite originating from the same RPF pool (**Extended Data 1b**). The differences seemed to be due to two technical effects. First, the ligation-based library included 29 nt reads that typically aligned with a mismatch at the 5′-most position, a known consequence of non-templated nucleotide addition to the 3′ end of cDNA after reverse transcription using retroviral RT (**Extended Data 1c**) [19]. These extended reads were essentially absent in the OTTR library. Second, the OTTR method begins with a terminal transferase reaction that adds an extra nucleotide to the RPF 3′ end to enable the template-jumping reverse transcription; computationally, this nucleotide is then trimmed off the sequenced read. If an RPF’s 3′ end is not extended but its 3′-most nucleotide is complementary to the adapter duplex primer’s 3′ overhang it can still be synthesized into the cDNA library, but consequentially, the 3′ nucleotide of the original RPF sequence will be trimmed. We are currently optimizing terminal transferase conditions to reduce this artifact. Despite these minor differences between OTTR and ligation-based libraries in the discrimination of RPF 5′ and 3′ ends, both methods captured the characteristic triplet periodicity of RPFs indicative of translation reading frames (**Figures 1d-e)**.

Technical factors such as ligase sequence preferences in ribosome profiling library generation have been shown to distort measurements of ribosome occupancy [17]. For example, the ligation-based library, but not the OTTR library, had a translatome-wide peak of read density for footprints with 5′ ends at the start codon, corresponding to an A-site assignment to the 5th codon after initiation (**Figures 1d-e**). We and others have noted this enrichment in many ribosome profiling data sets using ligation for cDNA library production [1, 19, 20]. The absence of this 5th-codon peak of read density in the OTTR library suggested that it could arise from a technical bias rather than an accumulation of ribosomes at this position in cells. Indeed, because all of these 5th codon RPFs have the same nucleotide sequence at their 5′ end, namely the AUG start codon, the sequence bias of enzymes used in the ligation-based protocol could explain their over-representation.

To quantify factors that influence RPF abundance in OTTR versus ligation-based libraries, we evaluated each library using iχnos, a neural network model of ribosome profiling data that predicts ribosome distribution along a transcript based on the sequences in a defined window encompassing the entire RPF [17]. Along with predicting the ribosome occupancy at each position, iχnos evaluates the predictive power of each feature it includes, namely, the sequences in and near the site of decoding. The identities of the codons in the A and P sites, which are paired with tRNAs within the ribosome, are expected to have the biggest impact on the speed of translation and thus the accumulation of RPFs. However, previous analysis with iχnos found that, in many RPF libraries made by ligation, the density of RPFs at a position was driven largely by the sequences at the 5′ and 3′ ends of the fragments, reflecting a strong influence of enzyme bias in RPF capture rather than the biology of ribosome decoding [17]. In keeping with this, models trained on our ligation-based RPF libraries showed the strongest effect from the identity of the sequences at the 5′ end of the RPF, confirming the strong bias introduced by CircLigase II (**Figure 1f**). Conversely, in OTTR libraries, we found the A-site and P-site codons contributed most to RPF abundance (**Figure 1f**). We conclude that the abundance of individual RPFs captured by OTTR was more indicative of tRNA-paired codon identities, and by inference, ribosome dwell time while decoding specific codons.

### P1 nuclease is a favorable alternative to RNase I for RPF production

*E. coli* RNase I has advantages in ribosome footprint generation, relative to alternatives such as combinations of RNases A and T1 or MNase, primarily due to a lack of sequence preference [16]. However, RNase I digestion is known to extensively fragment the rRNA, leading to two issues: 1) an accumulation of RPF-sized rRNA fragments that become overrepresented in the sequenced libraries; and 2) compromised structural integrity in a fraction of ribosomes, leading to subunit dissociation. We sought to utilize a sequence-independent RNase that would preferentially cleave unprotected mRNA while better sparing highly structured rRNA. The S1/P1 family of nucleases stood out for its general lack of nucleotide base specificity combined with strong preference for single-stranded RNA and production of cleavage products with 3′-terminal hydroxyls [21, 22]. Members of this family of nucleases bind single-stranded nucleic acids and cleave as endonucleases, producing either oligonucleotides or individual nucleotides when cleaving from a 3′ end. Although they are generally active at acidic pH, here we report P1 nuclease from *Penicillium citrinum* retains activity in a pH range compatible with ribosome stability in cell extracts [23].

To determine P1 nuclease digestion conditions suitable for ribosome profiling, we tracked the extent of inter-ribosome mRNA cleavage in crude cell lysates by measuring the conversion of polysomes to monosomes (**Extended Data 2a-b**). P1 nuclease digestion optimally separated yeast polysomes into monosomes when we lysed cells in standard pH 7.5 buffer then adjusted the pH of the lysate to 6.5 before adding P1 nuclease and incubating at 30 °C for one hour (**Extended Data 2a**). Compared to the RNase I digestion of an equivalent input sample, P1 nuclease digestion preserved more 80S monosomes and a small but reproducible amount of nuclease-resistant disomes (**Figure 2a**). P1 nuclease digestion did not produce free 40S and 60S subunits in excess of levels measured in the undigested control, whereas RNase I did increase the amount of free subunits, raising the possibility that it cleaved rRNA to the point of causing subunit dissociation (**Figure 2a**). We likewise found that P1 nuclease digestion of lysates from cultured human Calu-3 cells converted polysomes into monosomes with a small nuclease-resistant disome peak and minimal changes to free 40S and 60S subunits (**Figure 2b, Extended Data 2c**).

**Figure 2:**
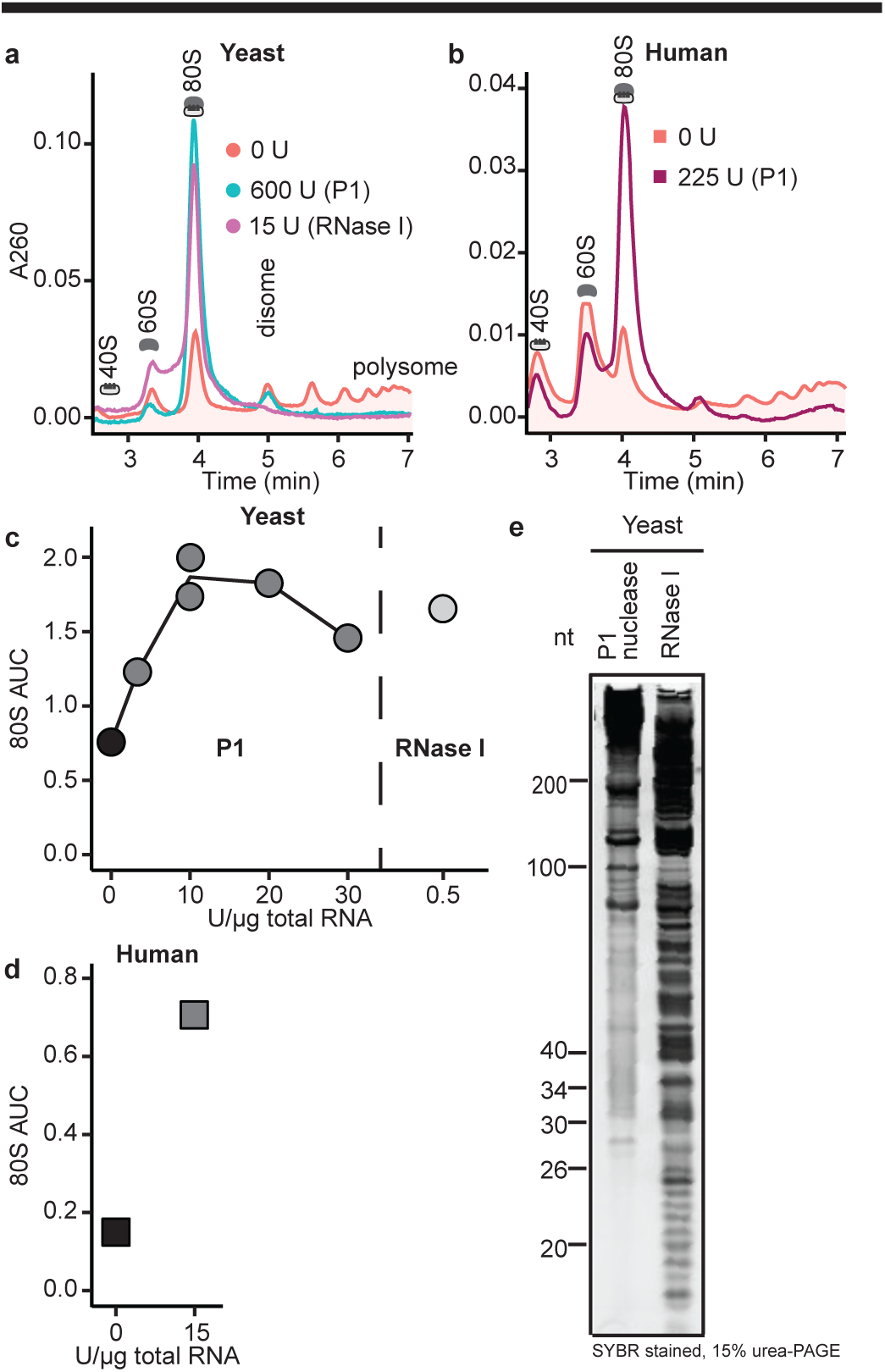
P1 nuclease collapse of polysomes into monosomes while limiting rRNA degradation. **a**, Sucrose density gradient polysome profiles of nuclease-treated yeast lysate, along with an undigested control. Each sample contained 30 µg of total RNA in 200 µL prior to nuclease digestion with the units (U) of enzyme indicated. P1 nuclease samples were adjusted to pH 6.5 prior to digestion and digested at 30°C for 1 hour. RNase I digestion was at room temperature for 45 min. **b**, As in (**a**), for human-derived Calu-3 cell lysate containing 15 µg of total RNA. **c**, Integrated monosome peak area from yeast lysate digested with various concentrations of P1 nuclease at 30 °C or RNase I at room temperature for an hour. Dark fill indicates undigested control, which was incubated without nuclease at 4 °C for an hour (n=1 for each condition, except n=2 for 10 U/µg). **d**, As in (**c**) for human Calu-3 cell lysate, but with P1 nuclease digestion at 37 °C. Dark fill indicates undigested control, which was incubated without nuclease at 4 °C for an hour (n=1 for each condition). **e**, Denaturing PAGE analysis of RNA extracted from ribosomal pellets after digestion with P1 nuclease or RNase I, as in (a), and stained with SYBR Gold.

Yeast lysate monosome yield, measured as the area under the curve (AUC) of the absorbance in the polysome profile’s 80S peak, reached an optimal plateau at 10-20 units (U) per µg of total RNA (300 U to 600 U of P1 nuclease in lysate containing 30 µg of RNA, **Figure 2c**). Using human lysate, 15 U/µg RNA likewise sufficed to produce monosomes (225 U of P1 nuclease in lysate containing 15 µg of RNA, **Figure 2d**), with optimal digestion at 37 °C rather than 30 °C (**Extended Data 2b-c**).

We next assessed rRNA integrity comparing RNase I and P1 nuclease digestion. We anticipated that conditions that favor RPF production over rRNA cleavage would result in less RNA within the RPF size range. RPFs alone have an estimated maximum yield of ∼4.5 ng RPFs per µg of total lysate RNA, assuming all ribosomes produce an RPF. After recovering ribosomes by ultracentrifugation, RNA was extracted from ribosome pellets and examined by denaturing PAGE and direct staining. Of note is the stark contrast in the RNA fragment profiles between RNase I and P1 nuclease digestion: RNase I generates a range of abundant small RNA fragment sizes that are heterogeneous in the range of RPF lengths, whereas P1 nuclease produces far less RNA in this size range (**Figure 2e**). This observation, coupled with the limited creation of free 40S and 60S subunits, suggests that P1 nuclease preferentially cleaves mRNA over rRNA relative to RNase I.

### P1 nuclease supports accurate ribosome profiling in yeast and human cells

To directly compare the RPFs produced by RNase I and P1 nuclease digestions, we generated pools of RNase I or P1 nuclease RPFs from the same yeast and human 293T cell lysates. We generated cDNA libraries using OTTR (**Extended Data 3a**) and sequenced them. P1 RPFs were longer than RNase I RPFs: P1 RPFs were 31 – 36 nt in yeast, compared to 26 – 29 nt RNase I RPFs, and in human cells P1 RPFs were 34 – 39 nt, compared to 28 – 32 nt for RNase I RPFs (**Extended Data 3b-c**). P1 RPF libraries yielded a larger fraction of reads mapping to mRNA (**Extended Data 3d-e**) and contained fewer discrete cytosolic rRNA fragment sequences of high abundance (**Extended Data 3f-g**). Of note, P1 nuclease produced more tRNA- and other non-coding (nc)RNA-derived reads relative to RNase I, particularly in human cells (**Extended Data 3d-e**). Mitochondrial rRNA-derived reads were detected in libraries made using both nucleases (**Extended Data 3h**). In OTTR RPF libraries made with RNase I or P1 nuclease, an increase in mean ribosome occupancy was detected near mRNA positions of translation initiation and termination for both yeast and human CDSs (**Figures 3a-b**). Gene-level ribosome occupancy measurements showed high correlation between nucleases for both yeast and human libraries (**Figures 3c-d**). P1 RPFs from initiating yeast and human ribosomes consistently had 5′ ends extended by 1 – 2 nt compared to RNase I footprints, positioning the A-site codon 16 – 17 nt from the 5′ end (**Figure 3e-f**). This increase in RPF length using P1 nuclease versus RNase I was also evident in RPFs from terminating yeast ribosomes: RNase I produced RPFs with 5′ ends located 14 – 15 nt upstream of the stop codon while P1 RPFs had 5′ ends 16 – 17 nt upstream (**Figure 3a**). Consistently, the heterogeneity of RPF lengths for any given ribosome A-site position primarily reflects a heterogeneous position of RPF 3′ end (**Figures 3e-f**).

**Figure 3:**
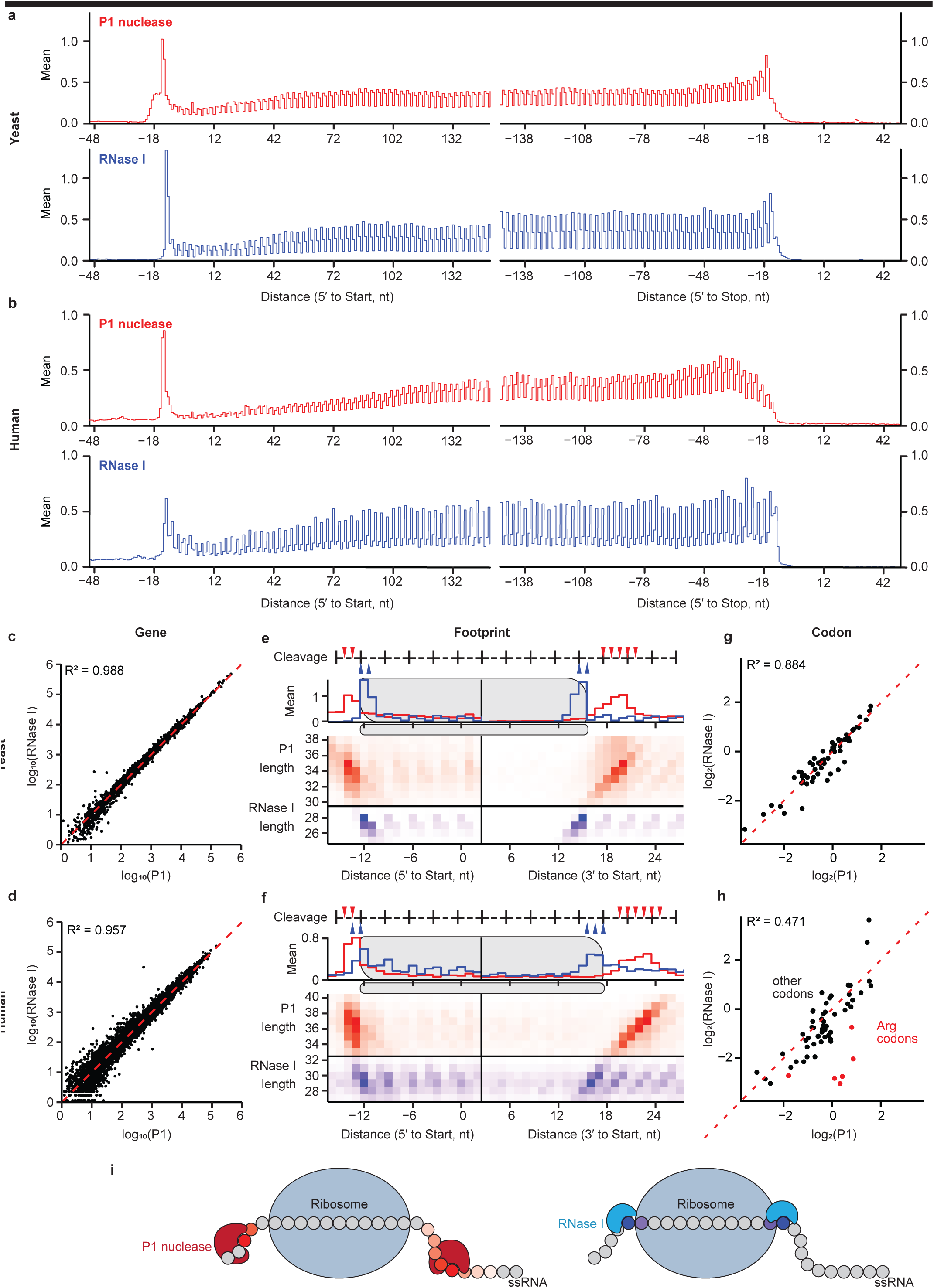
Highly similar ribosome profiling data from P1 nuclease or RNase I digestion. **a,** Metagene average profiles around the start (left) and stop (right) codons from yeast RPFs generated by P1 nuclease (top, red) or RNase I (bottom, blue) digestion. The 5′ ends of aligned reads were counted, and counts for each gene were rescaled by the mean codon count for the gene, excluding those RPFs that are aligned to the first 15 and last 10 codons, prior to averaging. **b**, As in (**a**), for human 293T cell RPFs generated by P1 nuclease (top, red) or RNase I (bottom, blue) digestion. **c**, Gene-level ribosome occupancy estimates from yeast RPFs generated by P1 nuclease and RNase I digestion, as in Fig. 1b. **d**, Gene-level estimates from human 293T cell RPFs generated by P1 nuclease and RNase I digestion. **e**, Average profile of yeast footprints at start codons for P1 nuclease (red, 30 – 40 nt) and RNase I (blue, 25 – 29 nt) libraries. Footprint alignments were counted separately for each gene monitoring read length as well as 5′ end position (left) and 3′ end position (right), then averaged as in (a). A heatmap shows footprint abundance according to length and end position (below), and the end position average summed across all lengths is shown (above each heatmap matrix. The 5′ and 3′ end averages are shown to the left and to the right, respectively, of a black vertical bar. A diagram of a translating ribosome footprint (top) indicates mRNA cleavage positions of P1 nuclease (red triangle) and RNase I (blue triangle). **f**, Average profile of human cell footprints at start codons for P1 nuclease (red, 33 – 40 nt) and RNase I (blue, 27 – 32 nt) libraries, as in (e). **g-h**, Comparison codon-level ribosome occupancy estimates from (**g**) yeast or (**h**) human cell RPFs generated by P1 nuclease and RNase I digestion, as in Fig. 1c. In (**h**), arginine codons are shown in red. **i**, Schematic of proposed P1 nuclease and RNase I cleavage sites around an mRNA-engaged ribosome. Increased frequency of an RPF terminal position is indicated by increasing color saturation.

We used the RPFs at initiating codons to empirically calibrate the offset between the 5′ ends of a footprint and the A- and P-site codons for both RNase I and P1 nuclease footprints. Across CDSs, inferred A-site codon occupancies correlated well for yeast RPFs produced by P1 nuclease and RNase I (R^2^ = 0.884) (**Figure 3g**). The correlation was somewhat lower between P1 nuclease and RNase I from human cell lysates (R^2^ = 0.471) driven primarily by an under-representation of arginine A-site codon assignments among RNase I RPFs (**Figure 3h**). In this study we selected RNase I RPFs of 26 – 34 nt, which would deplete the population of ∼21 nt RPF that arises from RNase I cleavage within an A-site when unoccupied by a tRNA [24, 25]. Codon-level differences between RNase I and P1 RPFs could reflect differences in empty A-site cleavage: RNase I cleaves within the A-site at slowly-decoded arginine codons and truncates the RPF, while P1 nuclease leaves the RPF in these ribosome complexes intact.

Indeed, several lines of evidence suggest that P1 nuclease is more sensitive to steric hindrance and thus less likely to cleave within the ribosome A-site. P1/S1 nuclease makes extensive contacts with several single-stranded nucleotides upstream of the cleaved phosphodiester bond [21, 26]. Moreover, P1 nuclease cleavage of DNA mismatches requires 2 – 3 nt of consecutive single-stranded nucleotides [27]. In the context of RPF generation, we propose that P1 nuclease requires several nucleotides of unobstructed mRNA in order to cleave. This requirement is satisfied for the mRNA exiting the ribosome, as evidenced by the uniform 5′ cleavage across a range of RPF lengths (**Figure 3i**). On the other side, mRNA upstream of the ribosome is more obstructed, giving rise to a wider range of possible cleavage positions, and thus, RPF lengths (**Figure 3i**).

### P1 nuclease preserves disome footprints from closely apposed ribosomes

In light of this proposed model for P1 nuclease cleavage, we wanted to know whether closely proximal ribosomes would interfere with P1 cleavage between ribosomes. Intriguingly, we noted the presence of a nuclease-resistant disome peak in the polysome profiles of P1 nuclease digested yeast and human cell lysates (**Figures 2a-b**). Adjacent ribosomes with minimal separation are known to arise when one ribosome stalls during elongation and a second, trailing ribosome collides with it. Such ribosome- ribosome collisions can trigger quality control pathways and broader cellular responses to translational stress such as suppression of translation initiation through phosphorylation of eukaryotic initiation factor 2 subunit α (eIF2α) [28, 29]. To test the hypothesis that P1 digestion preserves large RPFs spanning two collided ribosomes, we induced translational stalling at histidine codons by depriving yeast cells of charged histidyl-tRNA [30]. We induced the knock-down of histidyl-tRNA synthetase 1 (*HTS1*) mRNA in yeast that lacked Gcn2 and thus could not down-regulate translation initiation in response to uncharged tRNA accumulation (**Figure 4a**) [30, 31]. Knock-down of *HTS1* increased the disome population resolved by velocity sedimentation in a sucrose density gradient, following P1 nuclease digestion (**Figure 4b, Extended Data 4a**).

**Figure 4:**
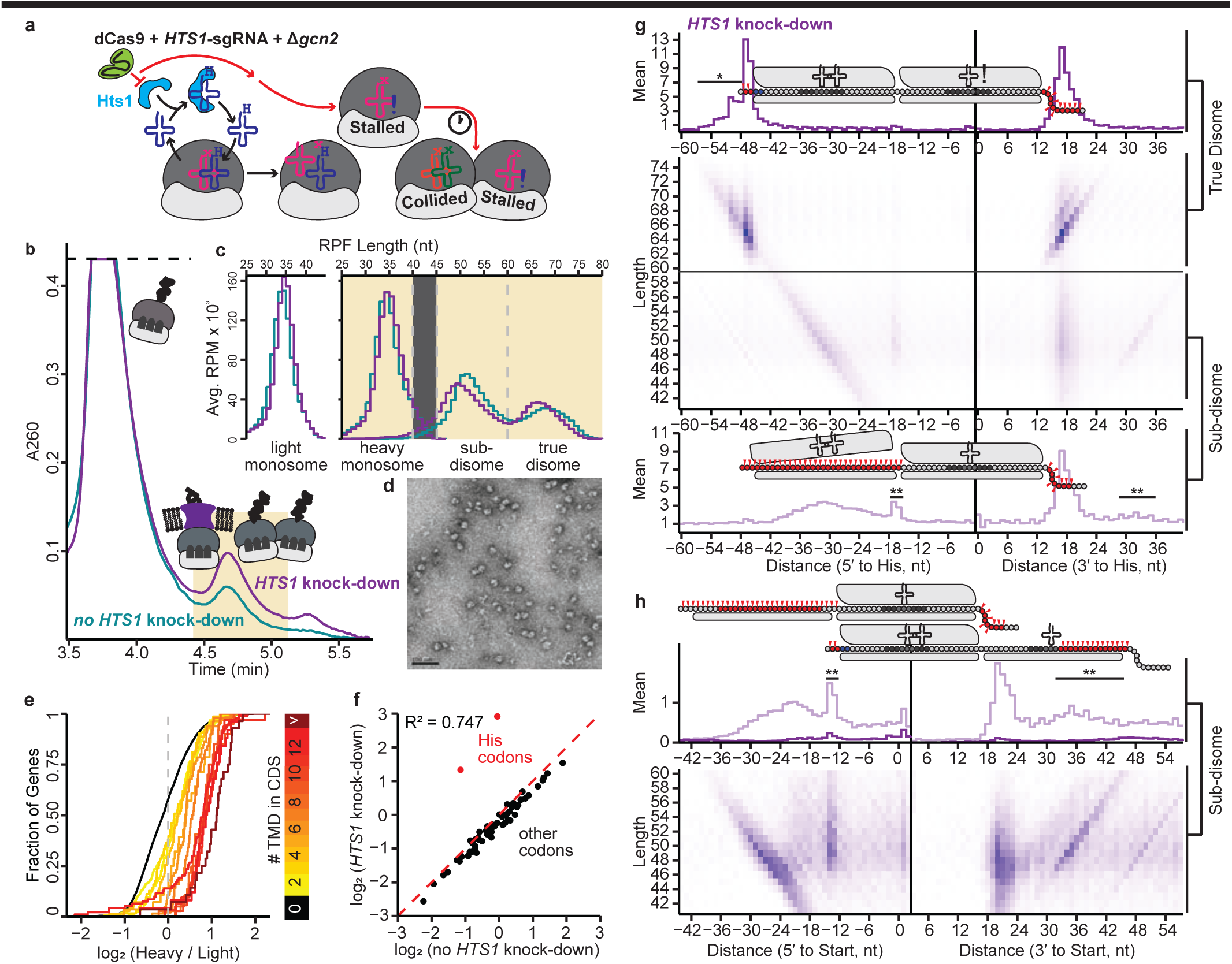
P1 nuclease disome and sub-disome footprints. **a**, Schematic of inducible, translatome-wide histidine codon stalling in yeast by depletion of histidyl-tRNA synthetase Hts1. **b**, Polysome profile of P1 nuclease digested yeast lysates with (purple) or without (teal) *HTS1* depletion. The yellow shaded box indicates the disome velocity sedimentation fraction collected. **c**, Read length distribution of RPF libraries from the monosome and disome velocity sedimentation peaks. The light monosome library (left) was generated from 30 – 40 nt RNA from the monosome fraction and the heavy monosome library was generated from 30 – 40 nt RNA in the disome fraction. The overall disome library was generated from 45 – 80 nt RNA in the disome fraction and computationally separated into sub-disome and true disome samples at the RPF length indicated by dashed line. The y-axis was represented in reads per million and averaged across the two replicates. **d**, Negative stain electron microscopy from a representative disome fraction from yeast lysate digested with P1 nuclease. Scale bar: 100 nm. **e**, Relative ribosome occupancy of genes comparing light and heavy monosome pools. Genes were stratified by the number of transmembrane domains they encode and cumulative distributions of occupancy between the heavy- and light-monosome libraries are plotted for each stratum. **f**, Comparison of codon-level ribosome occupancies between monosome samples with or without *HTS1* depletion. Histidine codons are shown in red, and Pearson’s R^2^ is reported for all codons including histidine. **g**, Average profile of yeast footprints in the true disome (top, 60 – 75 nt, purple line) and sub-disome (bottom, 40 – 59 nt, pink line) libraries around isolated histidine codons after *HTS1* depletion. The top schematic represents a ribosome-ribosome collision with the leading ribosome stalled at a histidine codon. The bottom schematic represents a similar ribosome collision, with the trailing ribosome undergoing RQC. The schematic 60S (top) and 40S (bottom) subunits with P- and A-site tRNAs are shown enclosing an mRNA (circles are nucleotides), with the E-, P-, and A-site nucleotides darkened. The A-site of the stalled ribosome was denoted with an exclamation point to symbolize the depletion of charged histidyl tRNA. Positions of P1 cleavage are denoted by red triangles. **h**, Average profile of yeast sub-disome footprints around start codons, as in (**g**). The overall occupancy of true disome footprints is also shown (dark purple histogram line). The schematic shows an 80S ribosome at initiation and an adjacent, upstream scanning pre-initiation complex (top) and an 80S ribosome at initiation and a downstream 40S subunit (bottom).

We isolated RNA from the monosome and disome velocity sedimentation fractions (**Figure 4b**). From the monosome peak we generated a library by size selection for 30 – 40 nt RNAs (**Figure 4c, Extended Data 4b**). In the disome fraction, negative-stain electron microscopy confirmed many ribosome pairs as well as potentially solitary ribosomes (**Figure 4d**). We therefore collected RPFs across size ranges from monosome to disome including a “heavy monosome” library of 30 – 40 nt RNA fragments (**Figure 4c, Extended Data 4b**). Sequencing of this heavy monosome library revealed RPFs similar to those in the conventional “light monosome” footprint library, but strongly enriched for transmembrane protein mRNAs (**Figure 4e**) [32]. We speculate that single ribosomes translating transmembrane proteins may sediment differently than other translating ribosomes due to their associations with protein translocation machinery or aggregation of nascent transmembrane regions [33].

Hts1 depletion induced dramatic ribosome stalling on histidine codons, which guided our expectations for RPF representation in the monosome and disome footprints. The average histidine codon occupancy in conventional monosome libraries was 5.3- and 7.3-fold higher for the CAU and CAC codons, respectively, when Hts1 was depleted (**Figure 4f**). We surveyed the distribution of RPFs at isolated histidine codons (i.e., no other histidine codon in a window from -30 to +10 codons; n=19,814) and noted a clear accumulation of footprint 5′ ends 16 – 17 nt upstream of the histidine codon with an RPF-length-dependent distribution of 3′ ends 14 – 21 nt downstream (**Extended Data 4c-d**). These RPF positions matched our inferred A-site offsets based on initiating ribosome (**Extended Data 4e-f**).

We also generated a library from 45 – 80 nt RNA fragments isolated from the disome peak (**Extended Data 4b**). Reads from this library fell into two size classes (**Figure 4c**). The larger fragments were roughly 65 nt, matching the footprint size anticipated for a pair of closely adjacent ribosomes. The smaller fragments were ∼50 nt, shorter than the ∼58 nt disome footprints produced by RNase I digestion [34] and thus probably not derived from two adjacent, intact 80S ribosomes. We computationally separated these RPFs into “true disome” footprints of at least 60 nt and “sub-disome” footprints shorter than 60 nt (**Figure 4c**).

Similar to the light monosome and heavy monosome RPFs, true disome RPFs also accumulated at histidine codons (**Figure 4g**). True disome RPFs had 5′ ends 46 – 47 nt upstream of the histidine codon and the typical length-dependent distribution of 3′ ends downstream (**Figure 4g**), as noted for monosome RPFs (**Extended Data 4c-d**). We also detected a small sub-population of true disome RPFs extended by 3 nt on their 5′ ends (**Figure 4g,** see asterisk). P1 nuclease digestion thus produced disome RPFs corresponding to a leading ribosome stalled on a histidine codon with a trailing ribosome in some cases 11 rather than 10 codons behind it.

The population of ∼50 nt sub-disome RPFs was unexpected. In most cases, sub-disome RPFs had 3′ ends 14 – 21 nt downstream of histidine codons, similar to monosome and true disome RPFs, but the 5′ ends were truncated 6 – 24 nt relative to true disome RPFs (**Figure 4g**). RNase I digestion is known to produce both a ∼58 nt true disome RPF and a 3′-truncated ∼51 nt RPF with cleavage at the unoccupied A-site of the leading stalled ribosome [34]. P1 nuclease did not appear to cleave within the unoccupied A-site of a stalled ribosome, however. Sub-disome RPFs with variable 5′ truncation but no 3′ truncation could arise from cleavage of mRNA partially protected at an intermediate stage of trailing (upstream) ribosome removal by RQC, with only the upstream 40S subunit stably engaged with mRNA (**Figure 4g,** depicted in the lower panel). Such intermediates might accumulate when RQC pathways are saturated by collisions following Hts1 depletion. We also noted a less abundant class of sub-disome RPFs with 5′ ends corresponding to monosomes stalled on the histidine codon and a broad distribution of 3′ ends downstream of this stall site. We hypothesize that these RPFs could correspond to stalled ribosomes with an adjacent, downstream 40S subunit (**Figure 4g,** RPF ends indicated by double-asterisks).

To extend disome analysis beyond positions of induced ribosome stalling at histidine codons, we examined the pattern of RPFs around start codons. Sub-disome RPFs were indeed enriched around start codons with very few true disomes (**Figure 4h,** the histogram line for true disomes is in dark purple). Similar to the profile at histidine-codon stalls, the majority of the sub-disomes at start codons had 5′ ends ranging from 18 – 40 nt upstream of the start codon and 3′ ends 18 – 24 nt downstream of the start codon, similar to the 3′ ends of RPFs corresponding to initiating monosomes (**Figure 4h**, **Extended Data 4e-f**). A minor population of sub-disome RPFs had 5′ ends 13 – 14 nt upstream of start codon, in common with initiating monosomes (**Extended Data 4e-f**), but a broader distribution of 3′ ends 32 – 45 nt downstream of the start codon (**Figure 4h,** see double-asterisks), paralleling the minor population of sub-disome RPFs at histidine codon stalls that have 5′ ends 17 – 16 nt upstream of a histidine codon (**Figure 4g,** see double-asterisks). Considering these results overall, we propose that P1 nuclease digestion preserves large RPFs, likely spanning an adjacent 40S subunit and 80S ribosome. These results highlight the utility of P1 nuclease digestion for mapping ribosome collisions and other higher-order ribosome complexes.

### P1 nuclease RPFs detect distinct ribosome configurations during short-ORF translation

In addition to ribosome stalling at histidine codons, Hts1 depletion, like other impairments to protein synthesis, triggers the integrated stress response through translational up-regulation of general control nondepressible 4 (*GCN4*) [35]. We recently reported that Hts1 depletion causes *GCN2*-independent induction of Gcn4 activity, and so we expected to see elevated *GCN4* translation in our data [30]. Indeed, we saw roughly 2-fold higher ribosome occupancy on the *GCN4* CDS after *HTS1* knock-down (**Figure 5a**) [36].

**Figure 5:**
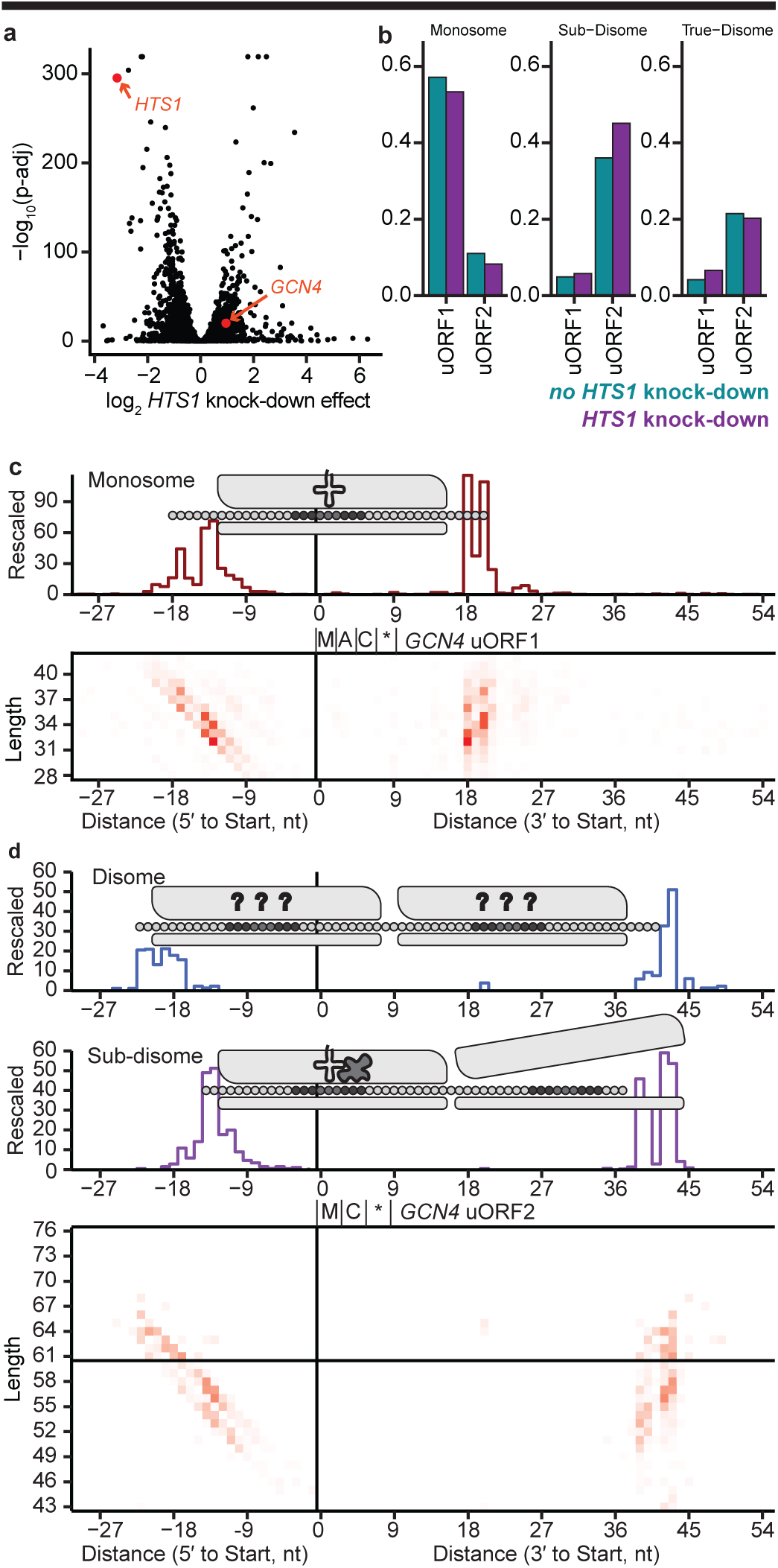
Monosome, sub-disome, and true disome occupancy profiles for *GCN4* uORF1 and uORF2. **a**, Differential expression analysis of monosome footprint occupancy across yeast genes, with versus without *HTS1* knock-down. *HTS1* and *GCN4* are shown in red. **b**, Average fraction of aligned footprints to uORF1 and uORF2 of *GCN4*. Counted alignments represented as fractions for each library before replicate libraries were averaged. Alignments were assigned to their respective ORF if both the 5′ and 3′ ends flanked the start and stop codons of the uORF but did not overlap another uORF. **c**, Monosome 5′ and 3′ profile of ribosomes at *GCN4* uORF1. Alignments were rescaled by the mean CDS codon occupancy of *GCN4* before replicates were summed together. **d**, Disome and sub-disome 5′ and 3′ profile of ribosomes at *GCN4* uORF2. Disome and sub-disome rescaled counts were normalized separately. Alignments for disome and sub-disome RPFs were rescaled by the respective mean codon count of *GCN4*’s CDS, excluding those RPFs that are aligned to the first 15 and last 10 codons, before replicates were combined by summing together.

Translational control of *GCN4* expression depends on short upstream ORFs (uORFs) in its 5′ leader. Previous ribosome profiling using RNase I suggested that most ribosome occupancy on the *GCN4* 5′ leader occurs over the first, 3-codon uORF (uORF1) [1]. The same was true for monosome-sized P1 nuclease RPFs with or without Hts1 depletion (**Figure 5b-c, Extended Data 5a-d**). Along with the typical monosome footprints for ribosomes initiating on uORF1, we observed a separate population of longer monosome footprints (36 – 39 nt) with 5′ ends 17 nt upstream of the start codon (**Figure 5c**). Upon *HTS1* knock-down, uORF1 monosome RPFs were slightly reduced while sub-disome and true disome sized footprints extending further upstream of the monosome RPFs were more apparent but not abundant (**Figure 5b, Extended Data 5a-d**), potentially explained by 43S queuing upstream of the initiating ribosome of uORF1 (see schematics in **Extended Data 5c-d**). In contrast to uORF1, sub-disome and true disome footprints were abundant at the second, 2-codon uORF (uORF2) under both conditions (**Figure 5b**, **Figure 5d, Extended Data 5e-h**). They included sub-disome footprints (50 – 52 nt) with 5′ ends 15 – 17 nt upstream of the terminating codon (**Figure 5d**) that were diminished after *HTS1* knock-down (**Extended Data 5e-h**). This unusual pattern of footprints suggests complex interactions between ribosomes at various stages of initiation, elongation, and termination on uORF2 (**Figure 5d, middle schematic**).

*CPA1* mRNA also harbors a regulatory uORF in its 5′ transcript leader. The 25-codon *CPA1* uORF is long enough to accommodate multiple ribosomes; it encodes the arginine attenuator peptide (AAP), which stalls ribosomes at the terminating codon when arginine is abundant [34, 37, 38]. Here, we observed many true disome RPFs produced when the leading ribosome was paused over the uORF stop codon (**Figure 6a**). Surprisingly, these footprints were longer and varied in length (67 – 75 nt) (**Extended Data 6a**) compared to collided disomes from ribosomes stalled on histidine codons after *HTS1* knock-down (63 – 65 nt) (**Figure 4g**). We were able to internally compare true disome footprints within the *CPA1* uORF because the last codon encodes histidine. *HTS1* knock-down shifted the 3′ ends of these true disome footprints one codon upstream, corresponding with the leading ribosome of each true disome now paused at the histidine codon one codon before termination (**Figure 6b, Extended Data 6b**). Notably, *HTS1* knock-down did not alter the 5′ end positions, only the footprint lengths and relative abundances of the individual true disomes footprints. This suggests the relative dwell times of trailing ribosomes had changed in response to the increase in leading ribosome stalling at the penultimate histidine codon rather than the termination codon. This led us to speculate that these longer true disome footprints could arise from nearly collided ribosomes separated by a gap of at least one codon.

**Figure 6:**
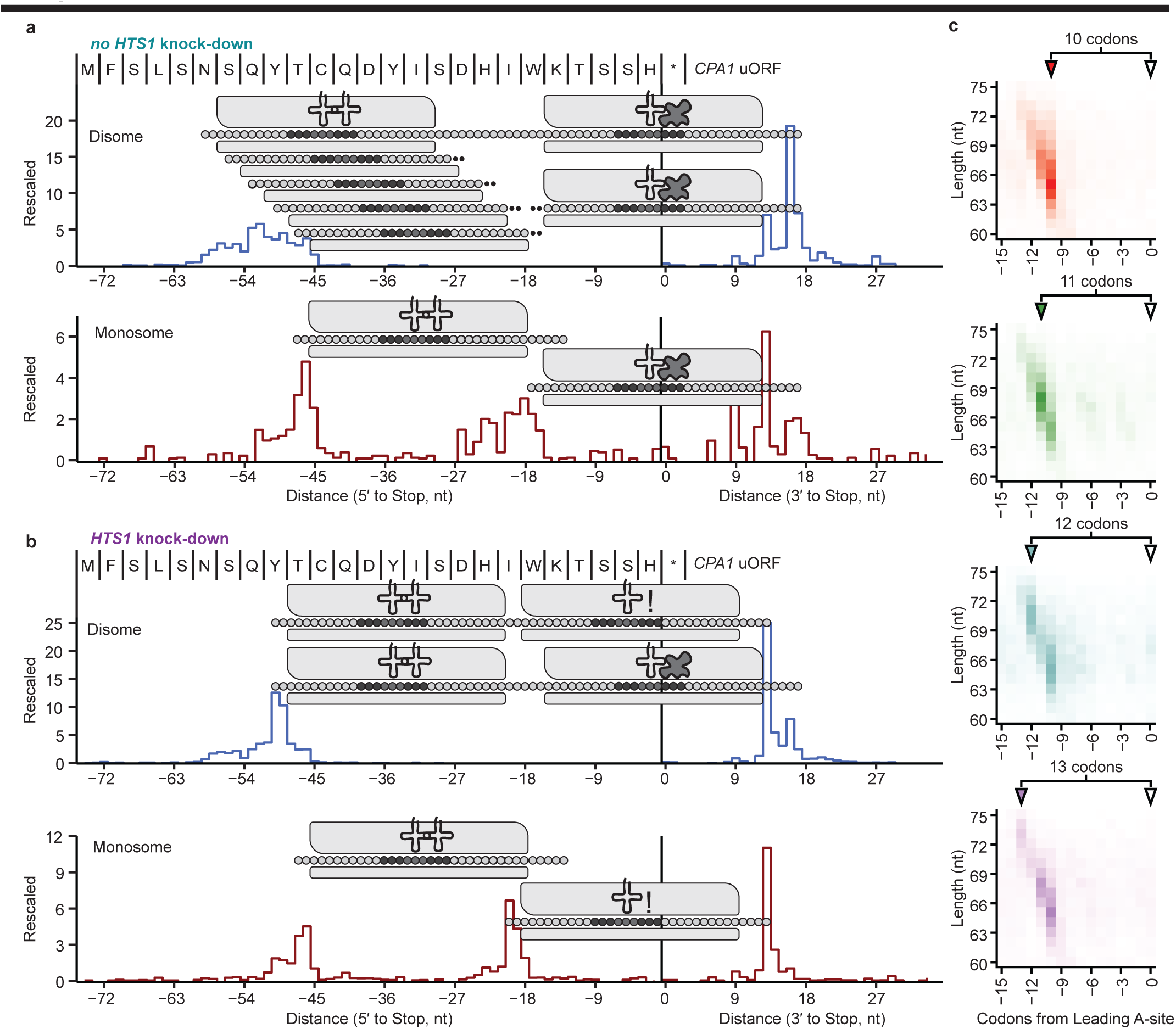
P1 nuclease captures nearly-collided pairs of ribosomes in both normal and Hts1-depleted conditions. **a**, Rescaled counts of disome (top) and monosome (bottom) 5′ and 3′ footprint ends with respect to the stop codon of the *CPA1* uORF in yeast without *HTS1* knock-down. Both a terminating ribosome (leading) and elongating ribosome (trailing) at various codon positions are schematized. **b**, As in (**a**) in yeast with *HTS1* knock-down. Both an elongating ribosome stalled at the penultimate histidine codon (leading) and an elongating ribosome (trailing) are schematized. **c**, Average rescaled counts of true disome footprints from *HTS1* knock-down yeast at CDS positions where pairs of histidine codons occur with 10 – 13 intervening non-histidine codons. The trailing ribosome A-site position is represented with respect to the leading ribosome stall at a histidine codon.

To investigate the P1 nuclease RPF signatures of nearly collided ribosomes on a translatome-wide basis, we examined ribosome occupancy after *HTS1* knock-down at positions where two histidine codons are separated by either 10, 11, 12, or 13 non-histidine codons. Histidine pairs separated by 10 codons yield the same ∼65 nt disome footprint that accumulates at isolated histidine codons (**Figure 6c,** top), consistent with a 10-codon gap as the minimal separation for a ribosome-ribosome collision. For codon gaps of 11, 12, and 13 we observed a steady increase in footprint lengths, principally from the 5′ ends, resulting in true disome footprints of ∼68, ∼71, and ∼74 nt, respectively (**Figure 6c**). We conclude that P1 nuclease can indeed protect pairs of nearly collided ribosomes with a gap of at least 1 – 3 codons between them, and we also establish that these nearly collided ribosomes accumulate under normal growth conditions in contexts such as termination on the *CPA1* uORF.

### Ribosome profiling without gel purification of RPFs

One remaining impediment to high-throughput ribosome profiling is the use of gel-based RPF size selection, which has been a necessity due to the high abundance and diversity of rRNA fragments produced by footprinting with RNase I. Given the relatively limited rRNA digestion by P1 nuclease (**Figure 2e**), we wanted to test whether the technically demanding gel-based RPF size selection could be avoided. We replaced gel-based RPF size selection with mirRICH small RNA enrichment, which relies on differential resuspension of small (< 200-300 nt) versus large RNAs from precipitated RNA [39]. We performed size selection only after OTTR cDNA synthesis, enriching library cDNAs with either monosome-size (30 – 45 nt) and/or disome-size (50 – 80 nt) inserts. We tested this approach using an independent P1 nuclease digestion of the *gcn2*Δ yeast lysates described above (**Figures 4 – 6**). After recovering ribosomes using a sucrose cushion and resuspending the pellet in polysome buffer and TRIzol, we split the material in half and recovered either total RNA or mirRICH small RNA as input for library synthesis (**Extended Data 7a**). OTTR cDNA synthesis reactions used ∼100 ng of total RNA or ∼40 ng of mirRICH-enriched RNA (∼1% of the total RNA or ∼10% of mirRICH RNA from the same amount of extract), and cDNA size selection was performed using OTTR cDNA generated from synthetic RNA input size markers (**Extended Data 7b**). The gene-level quantifications of ribosome occupancy using mirRICH or total RNA input libraries both matched results from libraries made with RPFs isolated by gel-based size selection (**Figure 7a, Extended Data 7c**). Using total RNA (**Extended Data 7c**), results for weakly expressed genes were distorted by low sequencing representation due to substantial presence of PCR duplicates (**Extended Data 7d**). This duplication arises because RPFs are relatively less abundant in total RNA than in the mirRICH-enriched RNA and therefore more cycles of PCR were required to make the sequencing library.

**Figure 7:**
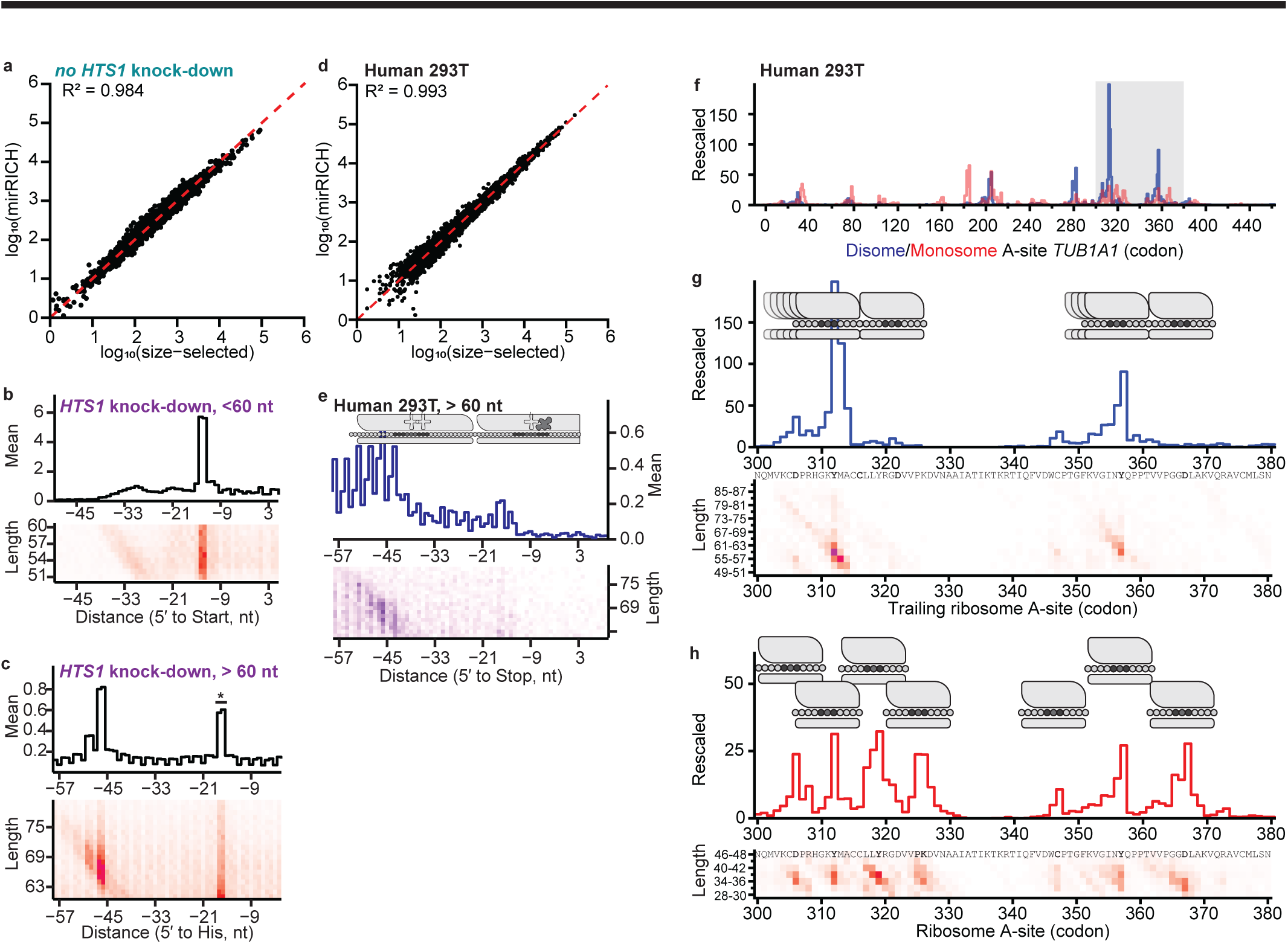
mirRICH and cDNA size-selection as alternatives to gel-based size selection of P1 nuclease RPFs. **a**, Gene-level estimates for CDS occupancy comparing mirRICH with light-monosome (i.e., size-selected) P1 RPF from yeast lysate without *HTS1* knock-down. mirRICH relied on cDNA size-selection whereas light-monosome relied on polysome fractionation followed by both RNA size-selection and cDNA size-selection. **b, c**, Footprint 5′ ends and length profile at initiation codons for (**b**) P1 sub-disome RPFs and (**c**) P1 true disome RPFs, captured by mirRICH from *HTS1* knock-down yeast lysate. The summed 5′ end profile is depicted at top and the contributions from each footprint length at each 5′ end are shown in the matrix at bottom. **d**, Gene-level estimates for CDS occupancy comparing mirRICH with gel purification size-selected P1 RPFs from human 293T cell lysates. **e,** Footprint 5′ ends and length profile at termination codons for P1 true disome RPFs captured by mirRICH from human 293T cell lysates. The summed 5′ end profile is depicted at top and the contributions from each footprint length at each 5′ end are shown in the matrix at bottom. **f**, Occupancy profile for monosomes (red) and trailing ribosome of the disome (blue) across the human *TUB1A1* ORF. The region in gray is expanded in **g**-**h**. **g, h**, Profile of A-site positions of (**g**) trailing ribosomes from true disome footprints and (**h**) monosome footprints, with length and position grouped by codon, from the gray region in (**f**). Monosomes (**h**) and disomes (**g**) are schematized above to emphasize differences in leading and trailing ribosome dwell time.

Next, we wanted to assess whether cDNA size selection of mirRICH libraries could enable bifurcated recovery of both monosome (30 – 45 nt) and disome (50 – 80 nt) sized libraries from the same OTTR cDNA library synthesis. We repeated P1 nuclease digestion of the *HTS1* knock-down yeast lysates (**Figures 4 – 6**), recovered ribosomes using a sucrose cushion, and isolated mirRICH-enriched RNA. We then carried out OTTR library construction and selected cDNA to individually isolate monosome- and disome-sized inserts. Promisingly, OTTR cDNA size selection was sufficient to separate disome-from monosome-sized reads (**Extended Data 7e**). Even without polysome fractionation of the input lysate, we captured the complexity of sub-disome footprints, such as 40S queuing upstream of initiating 80S ribosomes (**Figure 7b**), and true disome footprints from ribosome stalling at histidine codons (**Figure 7c**). A difference compared to libraries made from disomes purified by sucrose density gradients is the presence of some longer-than-monosome reads at mRNA positions expected to have stalled monosomes (**Figure 7c,** RPF ends indicated by asterisks). One possible explanation is that we captured highly extended monosome footprints where the P1 nuclease failed to fully degrade the unprotected mRNA 3′ of a ribosome.

We next applied mirRICH enrichment and cDNA size-selection to perform monosome and disome profiling from 293T human cell lysate. As in yeast, we performed bifurcated monosome and disome profiling using OTTR cDNA generated from mirRICH -enriched RNA purified from sucrose cushion pelleted ribosomes after P1 nuclease digestion (**Extended Data 7f**). We again observed that gel-purified and mirRICH monosome RFP library footprints yielded highly correlated measurements of ribosome occupancy (**Figure 7d**). We then examined libraries representing 50 – 80 nt RNAs, which should contain human disome and sub-disome footprints. Sub-disome RPFs accumulated around the sites of translation initiation, as noted in yeast, with few true-disome sized RPFs appearing before codons ∼15 – 18 (**Extended Data 7g**). At termination, we noted a dramatic enrichment of true-disome sized RPFs, consistent with a leading ribosome on the stop codon (**Figure 7e**). We also saw extended footprints at termination in these human cell lysates, likely reflecting extended footprints of nearly-collided ribosomes similar to those at the *CPA1* uORF termination (**Figure 7e**).

To investigate the global distribution of human disomes, we performed an analysis of disome-sized RPFs in well-translated CDSs to identify positions of trailing ribosome A-site codons, and grouped read lengths by codon as well (e.g., read lengths of 58, 59, and 60 nt mapped to 58 – 60 nt). With the assumption that human true disome RPFs are at least 58 nt, we determined the top five codons where disome accumulate to be in *RPS7* (codon Glu14), *TUBA1A* (Tyr312), *CCT5* (Arg24), *HSPE1* (STOP103), and *ACTB* (Arg147). For *TUBA1A,* which encodes α-tubulin, the major disome RPF was well separated from ribosomes at initiation or termination codons (**Figure 7f**). Two discrete sites of disome accumulation were evident, with trailing ribosomes at the codons for Tyr312 and Tyr357, along with extended RPFs from nearly-collided ribosomes (**Figure 7f-g**). At both sites of disome accumulation, monosome-sized footprints corresponding to the leading and trailing ribosomes of the disome were observed (**Figure 7h**). Notably, at a homologous position of yeast *TUB1* (amino acids 355 – 368 in *TUB1* corresponding to amino acids 354 – 367 in *TUBA1A*), slow kinetics of translation was observed associated with the recruitment of the TRiC/CCT chaperonin complex [40]. Disome accumulation at these positions could be explained by a similar phenomenon.

## Discussion

P1 nuclease digestion, mirRICH enrichment of RPFs, and OTTR cDNA library synthesis together provide a streamlined approach to ribosome profiling that offers many experimental advantages as well as increased biological insight from the preservation and capture of distinct classes of RPFs from closely approached and collided ribosomes. OTTR library generation provides adapter incorporation and reverse transcription simultaneously in a single-tube, 4-hour workflow, and as applied to ribosome profiling requires orders of magnitude less input RPF. As an additional benefit, OTTR RPF libraries show relatively low bias in capture of RNAs with different terminal nucleotides. We also demonstrate that P1 nuclease digestion improves monosome and disome yields relative to RNase I, reduces unwanted rRNA fragment contamination of the RPF RNA size range, and eliminates the risk of RNA degradation during RNase I RPF end-repair by phosphatase treatment. Footprints produced by P1 nuclease digestion are larger than those from RNase I, enabling improved precision of sequence mapping to the genome. In addition, OTTR protocol success using precipitation-based small RNA enrichment rather than more discriminating but labor-intensive gel-based size selection further simplifies RPF library generation. After OTTR cDNA synthesis, gel purification of Cy5-labeled products enables profiling of the monosome and disome footprints either together in a single library or separately. The simplicity, efficiency, and sensitivity achieved by combining P1 nuclease, mirRICH, OTTR, and Cy5-labeled cDNA eliminates days of labors and dozens of steps, providing the simplest ribosome profiling approach to date and suggesting a route towards automation. Moreover, these improvements may translate to a broader adoption of ribosome profiling over either mRNA sequencing or proteomic mass spectroscopy to access cell state.

Because P1 nuclease preserves higher-order ribosome complexes, ribosome profiling with P1 nuclease can offer additional insights about mechanisms governing translation initiation, elongation, and termination as well as different types of ribosome stalling, collision, and quality control. We did not detect depletion of RPFs by P1 nuclease cleavage of mRNAs within empty A-sites, in contrast to results from RNase I digestion [25, 41]. We did find that disomes resistant to P1 nuclease digestion include both collided ribosomes protecting a ∼65 nt footprint and larger disome footprints from nearby but not collided pairs of ribosomes. Lack of P1 nuclease cleavage in the gap between nearly-collided ribosomes provides an approach to investigate the occurrence and potential impact of closely approached ribosomes during their movement along an mRNA.

The sub-disome size class of RPFs preserved by P1 nuclease has not been previously investigated. To date we find these RPFs to be most abundant at start codons, stop codons, translation stalls, and uORFs. We suggest that they derive from proximity of an mRNA-engaged 40S subunit and 80S ribosome, with length heterogeneity at least in part arising from P1 nuclease cleavages spread across the mRNA region with 40S subunit only. At sites of 80S ribosome stalling, this 40S subunit may arise as a RQC intermediate, and distinctively in our study persist when RQC is saturated by translatome-wide delay in amino acid incorporation to nascent polypeptides. At translation start sites, the 40S may be a pre-initiation complex queued upstream of the initiating ribosome. While previous studies have proposed 40S-40S collisions, an initiation-specific RQC pathway involving pre-initiation complex collisions was described only recently [42, 43]. At termination, the pattern of sub-disome RPFs suggests that the 40S lies downstream of a terminating 80S, as could occur if an unrecycled small subunit was displaced into the 3′ untranslated region. The sub-disome and true disome RPFs generated by P1 nuclease hold promise for deeper insights into translation kinetics and ribosome-associated RNA and protein quality control.

## Acknowledgements

H.E.U., L.F., S.C.P., and K.C. were supported by NIH grants R35 GM130315 and DP1 HL156819, as well as the Bakar Fellows Program (to K.C.) and NIH Grant T32 GM007232 (to L.F.). N.T.I. was supported by NIH grant R01 GM130996. A.M. and L.F.L. were supported by the National Science Foundation under award number 1936069, by the National Institute of General Medical Sciences of the National Institutes of Health under award number R01GM132104 (to L.F.L.), and by a grant from the Rose Hills Foundation (to L.F.L.). We are thankful for the Vincent J. Coates Genomics Sequencing Laboratory QB3 Genomics, UC Berkeley, Berkeley, CA, RRID:SCR_022170, for sequencing support. We also thank the staff at the U.C. Berkeley Electron Microscope Laboratory for advice and assistance in electron microscopy sample preparation and data collection. We also wish to thank Ryan Muller, Sam Fernandez, Ryan Flynn, and past and present members of the Collins, Ingolia, and Lareau labs of U.C. Berkeley for their various levels of support in the development of this manuscript.

## Competing Interests

L.F, H.E.U., S.C.P, and K.C. are named inventors on patent applications filed by the University of California describing biochemical activities of RTs used for OTTR. H.E.U. and K.C. have equity in Karnateq, Inc., which licensed the technology and is producing kits for OTTR cDNA library preparation. N.T.I. declares equity in Tevard Biosciences and Velia Therapeutics.

## Materials and methods

### Polysome buffers for cell lysis and polysome profiling

For both yeast and human cell lysis, buffers and protocols described in McGlincy et al., 2017, were followed [19]. Briefly, polysome buffer used unless otherwise stated was 20 mM Tris-HCl pH 7.5, 150 mM NaCl, 5 mM MgCl2, and 1 mM DTT. If a pH 6.5 polysome buffer was required, 20 mM Bis-Tris pH 6.5 was used instead of Tris-HCl pH 7.5. If used for cell lysis the buffer included 1% (v/v) Triton X-100 and 25 U/mL Turbo DNase I. For sucrose density gradients, a polysome buffer was made with 10% or 50% (w/v) D-sucrose. For sucrose cushions, a polysome buffer was made with 1 M D-sucrose. Buffers were made fresh and stored on ice, and RNase-free commercial buffers were preferentially used, otherwise buffers were made in DEPC-treated water, filtered with a 0.22 µm pore, and when possible, autoclaved.

### Yeast culture and harvesting for polysome and ribosome profiling

For Figures 1 – 3 and Extended Data 1 – 3 BY4741 S288C wild-type yeast were used, but for Figures 4 – 6 and Extended Data 4 – 7 a yeast strain with constitutively expressed dCas9 and recombinantly deleted *GCN2* modified to inducibly express *HTS1*-sgRNA were used [30]. Unless otherwise stated, yeast cells were cultured in sterilized yeast extract and peptone media supplemented with 2% (v/v) dextrose (YEPD) at 30 °C. Yeast cells were back-diluted to an optical density (OD_600nm_) of 0.05 or lower and grown to OD_600nm_ of 0.5 – 0.6 in order to capture translation during sustained log-phase growth. As described in McGlincy et al., 2017, yeast cells were rapidly filtered, submerged in liquid nitrogen, and combined with polysome lysis buffer before the frozen mass was subjected to cryogrinding [19]. Pulverized yeast cells were thawed on ice and clarified by centrifugation at 20,000 x g for 30 minutes at 4 °C. The supernatant was aliquoted into pre-chilled tubes and snap-frozen in liquid nitrogen before long-term storage at -80 °C. Cycloheximide was not used in this study.

### Human adherent cell culture and harvesting for polysome and ribosome profiling

Human Calu-3 cell lysate was used in Figures 2 and Extended Data 2 while 293T cell lysate was used in Figure 3, Figure 6, Extended Data 3, and Extended Data 7. Cell cultures were grown in 15 cm dishes and harvested at least 24 hours after splitting with no more than 70-80% confluency. Cells were left undisturbed 12 hours before harvesting. At the moment of harvesting, cell media was rapidly aspirated and briefly replaced with 5 mL of ice-cold 1X phosphate buffered saline (PBS) before the culture dish was rested on a bed of lightly salted ice. PBS was thoroughly aspirated before 600 µL of polysome lysis buffer was added dropwise to the 15 cm culture dish. Cells were detached by cell scrapping and the lysate was moved to a pre-chilled tube. Lysates were left undisturbed for 10 minutes on a separate bed of unsalted ice to allow for DNase digestion. Lysates were gently pipetted with a 1 mL pipette tip to reduce viscosity before clarification by centrifugation at 20,000 x g for 30 minutes at 4 °C. The supernatant was aliquoted into pre-chilled tubes and snap-frozen in liquid nitrogen before long-term storage at -80 °C. Cycloheximide was not used in this study.

### RNA quantification of cell lysate

Through direct comparison with a standard curve of 0.0 – 1.0 ng/µL rRNA (ThermoFisher), the rRNA concentration of thawed lysates diluted in 1X TE buffer was quantified using the protocol for Quant-iT RiboGreen RNA kit (ThermoFisher). The assay volume was transferred to Minicell Borosilicate thin glass cuvette (Promega) and arbitrary fluorescence units were measured for each sample and the standard curve by the Glomax Multi Jr. Detection System (Promega) using the Fluorometer Blue module. A linear regression from the standard curve and dilution factor were used to estimate the RNA concentration in a given lysate.

### RNase I digestion of yeast and human lysate

*E. coli* RNase I (10 U/µL, Epicentre) digestion occurred at room temperature for 45 minutes while lysate gently rotated or mixed. Nuclease digestion of polysome buffered lysate was expressed in units of nuclease per µg of measured total lysate RNA (U/µg). Routinely, 0.5 U/µg of RNase I was sufficient for polysome collapse, and thus, ribosome profiling. In these experiments, we digested 30 µg of yeast lysate and 15 µg of Calu-3 or 293T cell lysate in a 200 – 300 µL volume. RNase I activity was quenched by placing lysate on ice. Digested lysates were either resolved on a sucrose density gradient or pelleted on a sucrose cushion. For ribosome profiling, the RNase I RPFs were size-selected from 26 – 34 nt after resolving on denaturing 15% urea-PAGE, as described [19].

### P1 nuclease digestion of yeast and human lysate

Optimal digestion by P1 nuclease (100 U/µL, New England Biolabs) required lysate to be adjusted to pH 6.5 before digestion. We found adding 7 µL of 300 mM Bis-Tris pH 6.0 per 100 µL of polysome buffered lysate at the point of nuclease digestion to be sufficient. Based on findings reported here, we routinely digest yeast lysate with 10 – 30 U/µg of P1 nuclease at 30 °C for 60 minutes, while we digested human cell lysate with 15 – 45 U/µg of P1 nuclease at 37 °C for 60 minutes. P1 nuclease activity was quenched by placing lysate on ice and returning the pH to 7.5. Digested lysates were either resolved on sucrose density gradients or pelleted on sucrose cushion. For ribosome profiling, the P1 RPFs were size-selected from 30 – 40 nt after resolving on denaturing 15% urea-PAGE.

### Polysome profiling

Sucrose density gradients were made in 14 x 89 mm Seton Open-Top Polyclear centrifuge tubes (Seton Scientific) from 6 mL of polysome buffered 10% (w/v) D-sucrose and 6 mL of polysome buffered 50% D-sucrose solution using the Gradient Master (BioComp Instruments). Using a wide-bore pipette tip, 300 µL of untreated or nuclease treated lysate was overlaid on top of the gradient. Lysate was resolved through the gradient by ultracentrifugation on a SW-41 rotor for 3 hours at 36,000 RPM in a 4 °C ultracentrifuge. The resulting polysome gradient was processed by the Gradient Master and analyzed at A_260nm_ using the BioRad Econo UV Monitor (BioRad) to measure the polysome profile. Polysome profiles which we quantitatively compared were either made from the same lysate for all conditions compared or all digest conditions for a given lysate were first normalized to its respective untreated lysate polysome profile before results were compared across conditions in different lysates. The AUC for various polysome features (40S and 60S, 80S, disome, polysome) were measured, and the ratio of 80S/polysome AUC was calculated for each polysome profile. We compared 80S/polysome AUC of a given condition to its respective untreated reference polysome profile to measure relative increase in 80S versus collapse of polysome for different digestions across lysates. When fractions were collected for downstream analysis, the collection apparatus was first slowly cleared by 5 mL of 3% H_2_O_2_ followed by 30 mL of nuclease-free water before resuming polysome fractionation. Fractions were timed so the monosome and disome peaks were collected in entirely separate fractions.

Monosome collection was from the trough after the 60S peak to the trough after the 80S peak. Disome collection followed next until we reached the trough between the disome and trisome peaks. Fractions were collected in pre-chilled tubes and snap-frozen in liquid nitrogen before storage at -80C. RNase-free reagents and filter-tip pipettes were used for each step.

### Sucrose-cushioned pelleting of ribosomes

To a 13 x 56 mm polycarbonate thick wall tube (Beckman Coulter), 200 to 300 µL of nuclease digested lysate was gently pipetted onto 900 µL of pre-chilled polysome buffered 1M D-sucrose using a wide-bore tip. Sample tubes were ultra-centrifuged for an hour at 100,000 RPM in a pre-chilled 4 °C TLA 110 rotor. The sucrose cushion was completely aspirated and the ribosome pellet was completely disrupted with a pipet tip before resuspend with 30 µL of 1X polysome buffer or nuclease-free water. After 10 minutes of incubation on ice, 300 µL of TRIzol reagent (ThermoFisher) was added, the entire volume was transferred to a pre-chilled tube, vigorously vortexed, and incubated for 10 minutes at room temperature. More TRIzol was added to any sample which remained turbid after incubation. Either Direct-Zol RNA purification, mirRICH, or long-term storage at -80 °C occurred next.

### Direct-Zol RNA extraction and Urea-PAGE RNA size selection of RNase I or P1 RPFs

Direct-zol RNA Purification (Zymo Research) was performed as described by the manufacturer, including the optional on-column DNase I digestion. RNA was eluted in 100 µL of nuclease free water and precipitated at -80 °C for ≥ 3 hours with 1 µL of 15 mg/mL GlycoBlue coprecipitant (ThermoFisher), 10 µL of 3 M sodium acetate pH 5.5, and 300 µL of 100% ethanol. The precipitate collected after centrifuging for 30 minutes at 20,000 x g at 4 °C was subsequently washed with 1 mL of -20 °C pre-chilled 70% ethanol, vortexed, incubated at -20 °C for 15 minutes, and centrifuged for 15 minutes at 20,000 x g at 4 °C. This step was performed twice. After the second wash, the RNA pellet was resuspended in 5 µL of 0.025% (w/v) bromophenol blue, 0.025% (w/v) xylene cyanol, 5 mM pH 8.0 EDTA, and 95% formamide (2X RNA loading dye) and either resolved by denaturing urea-PAGE electrophoresis or stored at -80 °C. We report precipitating the eluted Direct-Zol RNA and rigorous 70% ethanol washing as necessary steps to completely remove the salts carried over from TRIzol reagent before electrophoresis, which can interfere with accurate RNA size-selection. RNase-free reagents and filter-tip pipettes were used for each step.

Glassware and plasticware used for RNA gel electrophoresis were soaked in 10% bleach for 5 minutes before washed by COUNT-OFF™ Surface Cleaner (PerkinElmer) and liberally rinsing in double-distilled water. An RNase-free solution of 0.6X TBE, 7M UltraPure^TM^ urea (ThermoFisher), and 15% 19:1 acrylamide:bis-acrylamide (Ambion) was polymerized by adding 1% ammonium persulfate and 0.1% UltraPure^TM^ TEMED (ThermoFisher) and cast between borosilicate glass planes, allowing at least 30 minutes for polymerization. Once cast, gels were placed in a cleaned electrophoresis chamber in RNase-free 0.6X TBE running buffer. The wells were washed of excess urea, and the gel was pre-run at 150 V for 5 minutes. Prior to this, the precipitated RNA samples resuspended in 5 µL of 2X RNA loading dye were denatured at 80 °C for 90 seconds. Denatured RNA was kept on ice until loaded. Wells were washed again before 5 µL of denatured RNA was loaded. Wells were skipped between samples as necessary. When specified, a pair of either previously described RNase I RPF markers (RNA, 26 and 34 nt) [1] or a pair of P1 RPF markers (30 nt, 5′PO_4_-rNrGrA rCrArG rArCrU rGrArC rUrArC rUrCrA rCrArC rGrArA rCrArG rGrArN-3′OH; and 40 nt, 5′-PO_4_-rNrArG rUrCrG rUrCrU rCrArU rCrArG rGrUrC rUrCrU rCrArC rUrCrA rCrUrA rCrArC rArCrU rCrUrC rN-3′OH, Integrated DNA Technologies) were included. Electrophoresis was performed at 500 V for an hour. Gels were untouched after disassembly and manipulated exclusively by Saran-wrap or single-use blades to limit RNase and cross-nucleic acid contamination. Gels were stained for 1 minute with SYBR Gold (ThermoFisher). Gels were either analyzed on a Typhoon Trio (Cytiva) or Dark Reader Blue Light Transilluminator (Clare Chemical Research). RNase I RPFs were excised with the aid of the 26 nt to 34 nt maker while P1 RPFs were excised aided by the 30 nt to 40 nt marker. P1 disome RPFs were excised from 45 – 80 nt. We recommend excising 40 – 60 nt to select for the reported sub-disome P1 RPFs and 60 – 80 nt for reported true disome P1 RPFs.

### RNA elution from urea-PAGE

Several modifications were made to improve RNA elution by eliminating co-precipitates that inhibited OTTR cDNA synthesis. Firstly, we used an RNA elution buffer with 300 mM NaCl, 10 mM Tris-HCl pH 7.0, 1 mM EDTA, and 0.25% (v/v) SDS, which improved RNA stability during overnight room temperature elution while also reducing SDS co-precipitation when combined with ethanol. Secondly, gel slices were ground to mush using the tip of a pipette and the walls of a 1.5 mL tubes to maximize surface area. We routinely elute RNA overnight at room temperature in 400 µL of this RNA elution buffer. Gel bits are excluded from precipitation by centrifuging at 10,000 x g for 5 minutes at room temperature and removing only ∼350 µL of eluate. The eluate is combined with 1 µL of 15 mg/mL GlycoBlue coprecipitant and three volumes of 100% ethanol, and incubated for ≥ 3 hours at -80 °C. The RNA precipitate collected after centrifuging for 30 minutes at 20,000 x g at 4 °C was subsequently washed with 1 mL of -20 °C pre-chilled 70% ethanol, vortexed, incubated at -20 °C for 15 minutes, and centrifuged for 15 minutes at 20,000 x g at 4 °C. We routinely washed precipitates with 1 mL pre-chilled 70% ethanol 1 – 2 times. The 70% ethanol is completely removed and the pellet is allowed to airdry briefly before resuspending in 10 – 15 µL of RNase-free water. RNA concentration was estimated by NanoDrop. RNA was stored at -80 °C.

### RNase I RPF T4 polynucleotide kinase (PNK) dephosphorylation

For every 30 µg of treated lysate, we treated the RNase I RPFs with 5 U of T4 PNK (NEB, 10 U/µL) after RNA size selection [19]. We also routinely prepared 10X T4 PNK Buffer (700 mM Tris-HCl pH 7.5, 100 mM MgCl_2_, 50 mM DTT) with fresh DTT on the day of use. Anecdotally, we advise RNase I RPFs be first subjected to Oligo Clean and Concentrate (Zymo Research) purification before T4-PNK dephosphorylation.

### Monosome and disome profiling by P1 nuclease profiling

Polysome profiles fractions were collected for either negative stain electron microscopy or sequencing. For these experiments, lysate containing 150 µg of total RNA was *quantum satis* to 300 µL with pH 7.5 polysome buffer and supplemented with 21 uL of 300 mM Bis-Tris (pH 6.0) and 10 U/µg of P1 nuclease before incubating at 30 °C for 1 hour. Monosome and disome fractions were collected from polysome profiles as described above. Fractions were thawed on ice and concentrated using an Amicon Ultra-4 concentrator (Amicon). The single-use concentrator units were pre-washed with ice-cold polysome buffer and kept wet. Pooled fractions from the respective monosome and disome peaks were diluted to a volume of 4 mL in polysome buffer and then loaded onto a pre-washed concentrator unit and centrifuged for 5 min at 4,000 x g in a pre-chilled centrifuge. Each concentrator was reloaded with 4 mL of chilled polysome buffer and spun again for two cycles, and a final volume of 100 – 200 µL was recovered and either snap-frozen in liquid nitrogen and stored at -80 °C or immediately combined with 300 – 600 µL of TRIzol for Direct-Zol RNA extraction as described above. Extracted RNA was resolved by 15% denaturing urea-PAGE as described for pelleted ribosomes, and from the monosome peaks we collected RNA from 30 – 40 nt while from the disome peaks we collected both 45 – 80 nt and also 30 – 40nt. RNA was eluted and precipitated as described above.

### Δgcn2 *HTS1*-sgRNA inducible yeast culture, harvesting, and monosome and disome profiling

Cultures of Δgcn2 *dCas9-Mxi1::TetR::kanMXsyn P(RPR1)Tet-HTS1sgRNA::KlLEU2* yeast [30] were back-diluted from an overnight culture to an OD_600_ of 0.025 in either 750 mL YEPD with 250 ng/ml of anhydrotetracycline (for *HTS1* knock-down) or 250 mL of YEPD (i.e., no *HTS1* knock-down). The uninduced culture was harvested at an OD of 0.60, at which time we also harvested the induced culture at an OD_600_ of 0.21. Lysates were quantified as described, and 150 µg of lysate was diluted into 300 µL of polysome buffer and adjusted to pH ∼6.5 with 21 µL of 300 mM Bis-Tris pH 6. 1500 U of P1 was added to this pH adjusted lysate and incubated at 30 °C for an hour, as described above. Nuclease digestion and subsequent polysome profiling were carried out at least twice for each condition. The disome/80S AUC for each polysome profile was measured. The monosome and disome peaks were collected and concentrated with an Amicon Ultra-4 filter, RNA was extracted by Direct-Zol RNA, and either monosome-size (30 – 40 nt) or disome-size (30 – 40 nt, 45 – 60 nt) RPFs were selected from a denaturing 15% urea-PAGE. The RNA was eluted and quantified as described. Up to 40 ng of 30 – 40 nt RNA and up to 60 ng of 45 – 80 nt RNA was used for OTTR library generation. In total, there were two monosome, two disome, and two heavy monosome libraries generated for each condition.

### OTTR ribosome profiling library generation

OTTR library generation requires two recombinantly expressed, modified non-LTR retroelement RT proteins [**18, 44, Ferguson et al., in prep**]. Plasmids encoding RT proteins used in OTTR are available from AddGene: 2Bc-T MBP_BoMoC(ed)_6xH (JumpPol) and 2Bc-T MBP_BoMoC(ed)_F753A_6xH (TailPol). Detailed OTTR protocols and reagent preparation instructions are available upon request. To 40 ng of RNA in 9 µL of nuclease-free water, 2 µL of Buffer 1A (140 mM Tris-HCl pH 7.5, 1 M KCl, 14 mM DTT, 35% PEG-8000) and 1 µL of Buffer 1B (28 mM MnCl_2_, 3.5 mM ddATP) was added and mixed well before 1 µL of 10 µM TailPol was added. The reaction was incubated for an hour and 30 minutes at 30 °C before 1 µL of Buffer 1C (3.5 mM ddGTP) was added. The reaction was well mixed once more before incubating for an additional 30 minutes at 30 °C. Unincorporated ddRTP molecules were inactivated by adding 1 µL of Buffer 2 (80 mM MgCl_2_) and 1 µL of 1:1 rSAP (NEB):50% glycerol, mixing, incubating for 15 minutes at 37 °C. Then, 1 µL of Buffer 3 (100 mM EGTA), the reaction was mixed and incubated at 65 °C for 3 minutes. The reaction was immediately transferred to ice before 1 µL of Buffer 4A (10 mM MgCl2, 900 mM KCl, 40% PEG-6000, 4 mM dGTP, 0.8 mM dTTP, 0.8 mM dCTP, 0.04 mM dATP, 3 mM 2,6-diaminopurine-2′-deoxyribose-5′-triphosphate) and 1 µL of Buffer 4B (1.8 μM +1dY DNA/RNA adapter duplex primer, either 3.6 μM 3′C chimeric adapter template or 3.6 μM 3′CYN_5_ chimeric adapter template) was added and the reaction was vortexed. Then, 1 µL of 10 µM JumpPol was added and the reaction was well mixed before incubating at 37 °C for 30 minutes followed by a 70 °C incubation for 5 minutes. To aid complete cDNA recovery, 0.5 μg/μL RNase A (Sigma, R6513) was added to the reaction and it was incubated at 50 °C for 5 minutes. 80 µL of cDNA extraction buffer (50 mM Tris pH 8, 20 mM EDTA, 0.2% SDS) was added to the reaction followed by 100 µL of phenol:chloroform:isoamyl alcohol (25:24:1) (PCI); the tube was vortexed and the aqueous phase was recovered. RNA was then precipitated by adding 10 uL of 7.5 M ammonium acetate and 300 uL of 100% ethanol and then incubating at - 80 °C for 3 hours. For RNA templates > 40 nt, 0.5 U of Thermostable RNase H (NEB) was added to RNase A digestion, and Proteinase K was added to the cDNA extraction buffer and incubated at 50 °C for 15 minutes followed by denaturation at 95 °C for 5 minutes before PCI extraction.

### Ligation-based ribosome profiling library generation

The protocol from McGlincy NM et al., 2017 was followed for ligation-based ribosome profiling [19]. RNase I RPFs were dephosphorylated with T4 PNK before splitting samples for ligation-based and OTTR library generation.

### cDNA size-selection, elution, and quantification

OTTR reaction products were resuspended in bromophenol blue formamide loading dye (xylene cyanol was excluded as it co-migrates with the ∼75 nt primer dimer and interferes with Cy5 detection). The RNA oligonucleotide size markers used for RPF selection were carried through cDNA synthesis and used as positive controls and size markers for cDNA isolation. OTTR cDNAs were resolved by electrophoresis in 0.6X TBE 8% Urea-PAGE and visualized without staining using the fluorescence of the 5′ Cy5-dye. Excised urea-PAGE gel slices were crushed against the walls of a 1.5 mL tube using a sterile 1 mL pipette tip and cDNA was eluted by incubating at 70 °C for an hour in cDNA elution buffer containing 300 mM NaCl, 1 mM EDTA, and 10 mM Tris pH 8.0. cDNA was precipitated from the eluate using 3X volume of 100% ethanol followed by a 1 hour incubation at -80 C or snap-freezing in liquid nitrogen. The precipitate was centrifuged for 15 minutes at 20,000 x g at 4 °C and the cDNA pellet was washed at least once with 70% ethanol and resuspended in 25 µL double distilled, nuclease-free water. We followed a previously described procedure to quantify cDNA by real-time PCR in order to determine the number of PCR cycles needed for library amplification [19]. We typically used 10 – 12 PCR cycles, and diluted some cDNA libraries to ensure all libraries in a cross-comparison cohort (e.g., all libraries in Figure 3) were amplified with identical conditions.

### mirRICH small RNA enrichment from sucrose cushion pelleted ribosomes

The ribosome pellets were first completely resuspended in a 60 µl polysome buffer and then combined with 600 µL TRIzol and incubated for 10 minutes. The sample was then split in half, with 330 µL purified by Direct-Zol and 330 µl by mirRICH [39]. Direct-Zol RNA was eluted in 100 µL, 10 µL 3 M sodium acetate pH 5.5 and 300 µL of 100% ethanol was added, and RNA was precipitated at -80 °C for 3 hours. For mirRICH samples, we proceeded with TRIzol RNA extraction by adding chloroform, vortexing, centrifuging, and isolating the aqueous phase as described by the manufacturer. RNA was precipitated by adding 100% isopropanol and the mix was incubated on ice for 10 min. RNA was pelleted by centrifugation at 20,000 x g for 15 minutes at 4 °C but it was not washed with 75% ethanol. Instead, all liquid was removed carefully and the pellet was air dried for at least 2 hours. 10 µL of double-distilled water was added to the air-dried pellet and RNA was eluted for 10 minutes. The eluate was not mixed or vortexted during the elution period. 9 µL of the eluate liquid was removed, leaving the remaining pellet and ∼1 µL behind, and the eluate was further purified by RNA Clean & Concentrator-5 and quantified by Nanodrop.

### Ligation-based and OTTR adapter trimming

Ligation-based library reads were trimmed as described using Cutadapt (v3.2) [19]. A custom script was used to partition reads based on the ligated adapter’s linker sequence, allowing for up to one mismatch. For the OTTR library in Figure 1, the first base (usually a C or T) was removed from each read, and the DNA/RNA adapter duplex primer sequence (-a RGATCGGAAGAGCACACGTCTGAACTCCAGTCAC) was trimmed. For all libraries, any trimmed reads shorter than 20 or of poor quality (-q 10) were discarded. For OTTR libraries after Figure 1 the following Cutadapt pipeline was used to trim the read and annotate the UMI and TPRT base in the FASTQ ID: cutadapt -m 2 -a GATCGGAAGAGCACACGTCTGAACTCCAGTCAC $tmpvar.fastq | cutadapt -u 7 --rename=’{id} NTA={cut_prefix}’ - | cutadapt -u -1 -- rename=’{id}_{comment}_TPRT={cut_suffix}’ - | cutadapt -m 20 -q 10 -

### Yeast reference sequences

Yeast rRNA sequences RDN18-1, RDN18-2, RDN25-1, RDN25-2, RDN37-1, RDN37-2, RDN5-1, RDN5-2, RDN5-3, RDN5-4, RDN5-5, RDN5-6, RDN58-1, RDN58-2, Q0020, and Q0158 from SGD were concatenated and indexed by Bowtie. The yeast tRNA sequences and reference were prepared as described [45]. ncRNA reference was made from the ncRNA, snoRNA, and snRNA sequences from SGD and indexed by Bowtie.

Yeast gDNA from SGD was indexed by Bowtie. Two different mRNA references were used in this paper. For analysis related to iXnos we used the previously described mRNA reference, except we extended the 5′ untranslated region (UTR) and 3′ UTR sequences to include up to 50 bases [17]. We also constructed an mRNA reference where the SGD mRNA CDS sequence was matched with the longest annotated 5′ UTR and 3′ UTR sequence using a custom script that combines the CDS coordinates with the annotated BED coordinates [46, 47]. For both references, we confirmed these constructed sequences to be correct by both remapping them to the genomic reference and confirming the expected polypeptide can be translated from the sequence. In total, 5432 yeast mRNA sequences were constructed with either an annotated 5′ UTR and/or 3′ UTR sequence. 432 Ty-element mRNA sequences and 396 intron-containing, CDS-only, or frameshifting sequences were identified and isolated to a separate reference

### Human reference sequences

Human rRNA sequences NR_023379.1, NR_146151.1, NR_146144.1, NR_145819.1, NR_146117.1, NR_003285.3, NR_003286.4, X12811.1, and NR_003287.4 from NCBI and ENST00000389680.2 and ENST00000387347.2 from Ensembl were concatenated and indexed by Bowtie. The human tRNA sequences and references were prepared as described [45]. ncRNA from Ensembl and lncRNA from Gencode were concatenated and indexed by Bowtie. The primary assembly of the human gDNA from NCBI, GRCh38, was indexed by Bowtie. 18,640 NCBI RefSeq MANE v0.95 transcripts were indexed by Bowtie, and the 5′ UTR, CDS, and 3′ UTR lengths were parsed from the GenBank entries using a custom script. The 13 mitochondrial mRNA sequences from Ensembl were also included in the mRNA reference, but typically excluded from analysis.

### General ribosome profiling bioinformatics

For both human and yeast, we used a sequential mapping pipeline to remove contaminating reads from RPFs. Trimmed reads were removed by either mapping to the rRNA reference (bowtie -v 3 -a --best --norc), mapping to the tRNA reference by tRAX [45], mapping to the ncRNA reference (bowtie -v 3 -a --best --norc), or by mapping to the mtDNA reference (bowtie -v 2 -a --best --norc). The remaining reads were then mapped to the mRNA references (bowtie -v 2 -m 200 -a --norc) and sorted (samtools sort -n). Lastly, we sequentially mapped remaining reads to the gDNA reference (bowtie -v 2 -a --best) to identify any remaining biologically-derived reads before mapping unaligned reads to a reference of size-selecting oligos and adapter oligos (bowtie -v 2 - a --best) to flag any remaining read as either technical artifacts or unmapped.

To quantify gene-level CDS occupancy from multi- and single-mapping alignments we first used Bedtools intersect (v2.25.0) to exclude reads which aligned outside of the +15th to -10th codon of the CDS, excluding either 5′ UTR or 3′ UTR mapping reads or reads near either initiating or terminating ribosome, which do not behave like elongating ribosomes. Multi-mapping reads were handled by RSEM (rsem-calculate-expression --alignments --strandedness forward --seed-length 20 --sampling-for-bam), and the resulting tables of counted CDS alignments per gene were normalized and analyzed by DESeq2 [31763789]. Two replicates were produced per condition, except for Figure 1B’s which were a direct comparison of two library methods from a single RPF sample.

### Initiation, termination, and solitary codon occupancy profiles

Only genes with ≥ 50 nt of annotated 5′ UTR, ≥ 450 nt of annotated CDS, ≥ 50 nt of annotated 3′ UTR, and ≥ 1 reads per codon were considered for this analysis.

Alignments were counted based on read length and either 5′ or 3′ position relative to the first base of a codon of interest (i.e., either start codon, stop codon, or solitary codon), and rescaled by average number of reads per codon for a given gene. Rescaled counts for each read length and relative position were summed and divided by the either the number of genes under consideration (i.e., for features that exist once per gene, like initiation and termination) or the number of windows under consideration (i.e., for solitary histidine codon loci across the translatome).

### A-site occupancy comparisons

Only cytosolic genes with ≥ 50 nt of annotated 5′ UTR, ≥ 450 nt of annotated CDS, ≥ 50 nt of annotated 3′ UTR, and ≥ 1 reads per codon were considered for this analysis.

Alignments outside of the first 15 and final 10 codons were ignored. A-site offsets were predetermined based on alignments to the initiation profile. A-site assignments were counted for each gene and rescaled by average number of reads per codon. The rescaled occupancy for each codon, excluding either initiation or termination codons, were averaged across the translatome, and the final A-site occupancy estimates were rescaled out of 60 before comparing.

### iχnos full-model and leave-one-out models

iχnos provides an example transcriptome reference for yeast. We extended the 5′ UTR and 3′ UTR sequences of the yeast transcriptome to include a total of 50 nt of both 5′ UTR and 3′ UTR sequences. After rRNA-, tRNA-, and ncRNA-mapping reads were excluded, Bowtie was used to map the reads against iχnos’s yeast mRNA transcriptome (bowtie -v 2 --norc -a -m 250), and multi-mapping reads were reconciled by RSEM (rsem-calculate-expression --alignments --strandedness forward --seed-length 20 -- sampling-for-bam). RSEM handles multi-mapping reads by assigning a weight to an alignment based on how well it scored in RSEM. iχnos can use RSEM weights, but can also use the highest-scoring multi-mapping read, which is what we used here.

A total of 1000 genes which met the criteria to be studied by iχnos were considered across each library evaluated in this study. These genes were split into training (n=674) and testing (n=326) gene sets. iχnos was run with paralog-sensitive parameters as recommended. For each library, the RPF alignments ranging from 25 to 29 nt in lengths were selected, and those which mapped to codons after the 20th but before the -20th codon (elongating-only) were evaluated. The 5′ alignments were collapsed to the codon-level using A-site offset, and the codon-level counts for a given gene were normalized by the gene’s average occupancy, providing a mean-occupancy scale for each codon. From this, we used a previously described iχnos model-design which explores a sequence space of 13 codons from the -5th codon upstream to the +7th codon downstream (n5p7) of a given A-site. We first trained the n5p7-model five times for 45 epochs at a learning rate of 16 for each library, measuring Pearson’s correlation of the predicted occupancy for the test-genes versus actual occupancy for each replicated trained-model. We subsequently performed the same training regimen on 13 different model designs, except this time we only evaluated 12 of the 13 codons, leaving one out (LOO) for each of the 13 different models. To interpret which codon had the most impact on the model’s predictions, we measured the average change in Pearson’s correlations for each of the 13 LOO-models compared to the full-model.

## Data availability

High-throughput sequencing data has been deposited with the NCBI Short Read Archive. Long-read sequencing linking tethering constructs and barcodes is available at SRR10355648, and short-read sequencing quantifying the barcodes is available at SRR10353306 through SRR10353315, as described below.

Acession; LibraryName; Title SAMN32928056; McGlincy; Ligation based ribosome profiling from a pool of yeast RNase I digested RPFs (technical replicate 1 with linker CGTAA, technical replicate 2 with linker GCATA was not successful).

SAMN32928057; OTTR1; OTTR ribosome profiling from a pool of yeast RNase I digested RPFs (technical replicate 1, paired with McGlincy linker CGTAA).

SAMN32928058; OTTR2; OTTR ribosome profiling from a pool of yeast RNase I digested RPFs (technical replicate 2, paired with McGlincy linker GCATA).

SAMN32928059; yP1; Nuclease digestion replicate 1 of yeast lysate with P1 nuclease.

SAMN32928060; yP2; Nuclease digestion replicate 2 of yeast lysate with P1 nuclease.

SAMN32928061; yR1; Nuclease digestion replicate 1 of yeast lysate with RNase I.

SAMN32928062; yR2; Nuclease digestion replicate 2 of yeast lysate with RNase I.

SAMN32928153; hP1; Nuclease digestion of biological replicate 1 of 293T cell lysate with P1 nuclease.

SAMN32928154; hP2; Nuclease digestion of biological replicate 2 of 293T cell lysate with P1 nuclease.

SAMN32928155; hR1; Nuclease digestion of biological replicate 1 of 293T cell lysate with RNase I.

SAMN32928156; hR2; Nuclease digestion of biological replicate 2 of 293T cell lysate with RNase I.

SAMN32928063; M2; Monosome-sized RPFs from monosome fraction of a polysome profile after P1 nuclease digestion, biological replicate 1 of untreated cells (no HTS1 knock-down).

SAMN32928064; M3; Monosome-sized RPFs from monosome fraction of a polysome profile after P1 nuclease digestion, biological replicate 2 of untreated cells (no HTS1 knock-down).

SAMN32928065; M4; Monosome-sized RPFs from monosome fraction of a polysome profile after P1 nuclease digestion, biological replicate 1 of anhydrotetracycline treated cells (HTS1 knock-down).

SAMN32928066; M5; Monosome-sized RPFs from monosome fraction of a polysome profile after P1 nuclease digestion, biological replicate 2 of anhydrotetracycline treated cells (HTS1 knock-down).

SAMN32928067; DM2; Monosome-sized RPFs from disome fraction of a polysome profile after P1 nuclease digestion, biological replicate 1 of untreated cells (no HTS1 knock-down).

SAMN32928068; DM3; Monosome-sized RPFs from disome fraction of a polysome profile after P1 nuclease digestion, biological replicate 2 of untreated cells (no HTS1 knock-down).

SAMN32928069; DM4; Monosome-sized RPFs from disome fraction of a polysome profile after P1 nuclease digestion, biological replicate 1 of anhydrotetracycline treated cells (HTS1 knock-down).

SAMN32928070; DM5; Monosome-sized RPFs from disome fraction of a polysome profile after P1 nuclease digestion, biological replicate 2 of anhydrotetracycline treated cells (HTS1 knock-down).

SAMN32928071; D2; Disome-sized RPFs from disome fraction of a polysome profile after P1 nuclease digestion, biological replicate 1 of untreated cells (no HTS1 knock-down).

SAMN32928072; D3; Disome-sized RPFs from disome fraction of a polysome profile after P1 nuclease digestion, biological replicate 2 of untreated cells (no HTS1 knock-down).

SAMN32928073; D4; Disome-sized RPFs from disome fraction of a polysome profile after P1 nuclease digestion, biological replicate 1 of anhydrotetracycline treated cells (HTS1 knock-down).

SAMN32928074; D5; Disome-sized RPFs from disome fraction of a polysome profile after P1 nuclease digestion, biological replicate 2 of anhydrotetracycline treated cells (HTS1 knock-down).

SAMN32928075; TetMMRMonoR1; mirRICH small RNA enrichment and cDNA size-selection of P1 nuclease digested RPFs from a sucrose cushion pellet, biological replicate 1 of untreated cells (no HTS1 knock-down).

SAMN32928076; TetMMRMonoR2; mirRICH small RNA enrichment and cDNA size-selection of P1 nuclease digested RPFs from a sucrose cushion pellet, biological replicate 2 of untreated cells (no HTS1 knock-down).

SAMN32928077; TetMDZMonoR1; Direct-Zol RNA purification and cDNA size-selection of P1 nuclease digested RPFs from a sucrose cushion pellet, biological replicate 1 of untreated cells (no HTS1 knock-down).

SAMN32928078; TetMDZMonoR2; Direct-Zol RNA purification and cDNA size-selection of P1 nuclease digested RPFs from a sucrose cushion pellet, biological replicate 2 of untreated cells (no HTS1 knock-down).

SAMN32928079; TetPMRMonoR1; mirRICH small RNA enrichment and cDNA size-selection of P1 nuclease digested RPFs from a sucrose cushion pellet, biological replicate 1 of anhydrotetracycline treated cells (HTS1 knock-down).

SAMN32928080; TetPMRMonoR2; mirRICH small RNA enrichment and cDNA size-selection of P1 nuclease digested RPFs from a sucrose cushion pellet, biological replicate 2 of anhydrotetracycline treated cells (HTS1 knock-down).

SAMN32928081; TetPMRDiR1; mirRICH small RNA enrichment and cDNA size-selection of P1 nuclease digested disome-sized RPFs from a sucrose cushion pellet, biological replicate 1 of anhydrotetracycline treated cells (HTS1 knock-down).

SAMN32928082; TetPMRDiR2; mirRICH small RNA enrichment and cDNA size-selection of P1 nuclease digested disome-sized RPFs from a sucrose cushion pellet, biological replicate 2 of anhydrotetracycline treated cells (HTS1 knock-down).

SAMN32928157; T293MRMonoR1; mirRICH small RNA enrichment and cDNA size-selection of P1 nuclease digested RPFs from a sucrose cushion pellet, biological replicate 1 of 293T cell lysate (different batch of lysates separate from hP1, hP2, hR1, hR2).

SAMN32928158; T293MRMonoR2; mirRICH small RNA enrichment and cDNA size-selection of P1 nuclease digested RPFs from a sucrose cushion pellet, biological replicate 2 of 293T cell lysate (different batch of lysates separate from hP1, hP2, hR1, hR2).

SAMN32928159; T293MRDiR1; mirRICH small RNA enrichment and cDNA size-selection of P1 nuclease digested disome-sized RPFs from a sucrose cushion pellet, biological replicate 1 of 293T cell lysate (different batch of lysates separate from hP1, hP2, hR1, hR2).

SAMN32928160; T293MRDiR2; mirRICH small RNA enrichment and cDNA size-selection of P1 nuclease digested disome-sized RPFs from a sucrose cushion pellet, biological replicate 2 of 293T cell lysate (different batch of lysates separate from hP1, hP2, hR1, hR2).

SAMN32928161; T293SSMonoR1; RNA size-selection and cDNA size-selection of P1 nuclease digested RPFs from a sucrose cushion pellet, biological replicate 1 of 293T cell lysate (different batch of lysates separate from hP1, hP2, hR1, hR2).

SAMN32928162; T293SSMonoR2; RNA size-selection and cDNA size-selection of P1 nuclease digested RPFs from a sucrose cushion pellet, biological replicate 1 of 293T cell lysate (different batch of lysates separate from hP1, hP2, hR1, hR2).

Publically available data sets used here include *S. cerevisiae* genome annotations from http://sgd-archive.yeastgenome.org/sequence/S288C_reference/genome_releases/S288C_reference_genome_R64-2-1_20150113.tgz

## Code availability

Custom software and a workflow used to analyze data in this study are provided in Zenodo at https://zenodo.org/record/7574339#.Y9MrtHbMKrQ

---

**Extended Data Figure 1:**
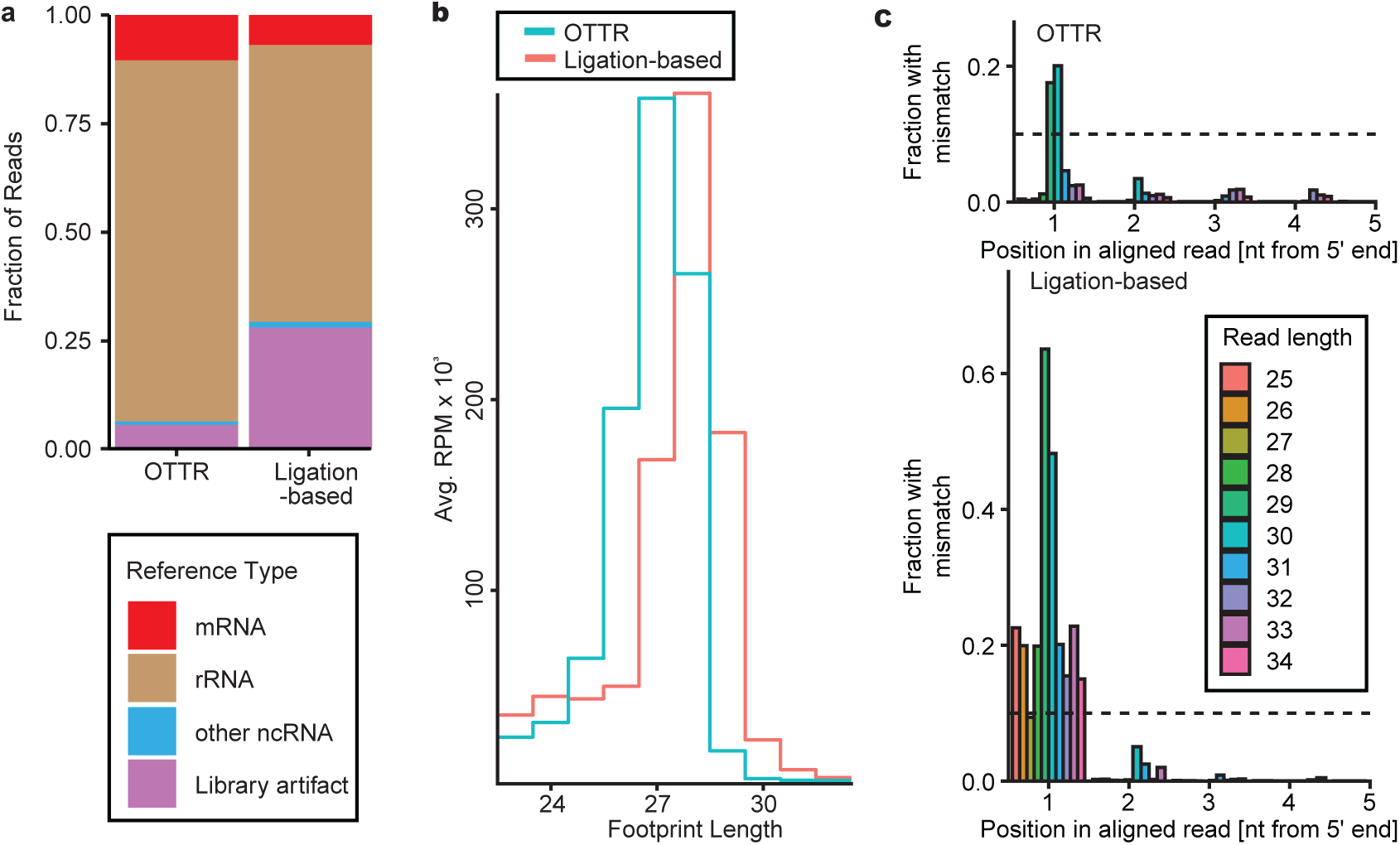
Additional comparison of RNase I RPF libraries by ligation-based or OTTR protocol. **a**, Fraction of RNase I RPF cDNA library sequencing reads mapped to each transcript class. Library generation artifacts included sequences that were adapter-only, shorter than 15 bases, or unmapped. **b**, Read length distribution of OTTR (blue) and ligation-based (red) RNase I RPFs from the CDS, excluding those RPFs that are aligned to the first 15 and last 10 codons. Counts were represented in RPM and averaged across replicates **c**, Fraction of reads with a mismatch near the 5′ end of the footprint from CDS aligned RPFs, stratified by read length. For this analysis in particular, alignments were permitted to only have a single mismatch to the reference.

**Extended Data Figure 2:**
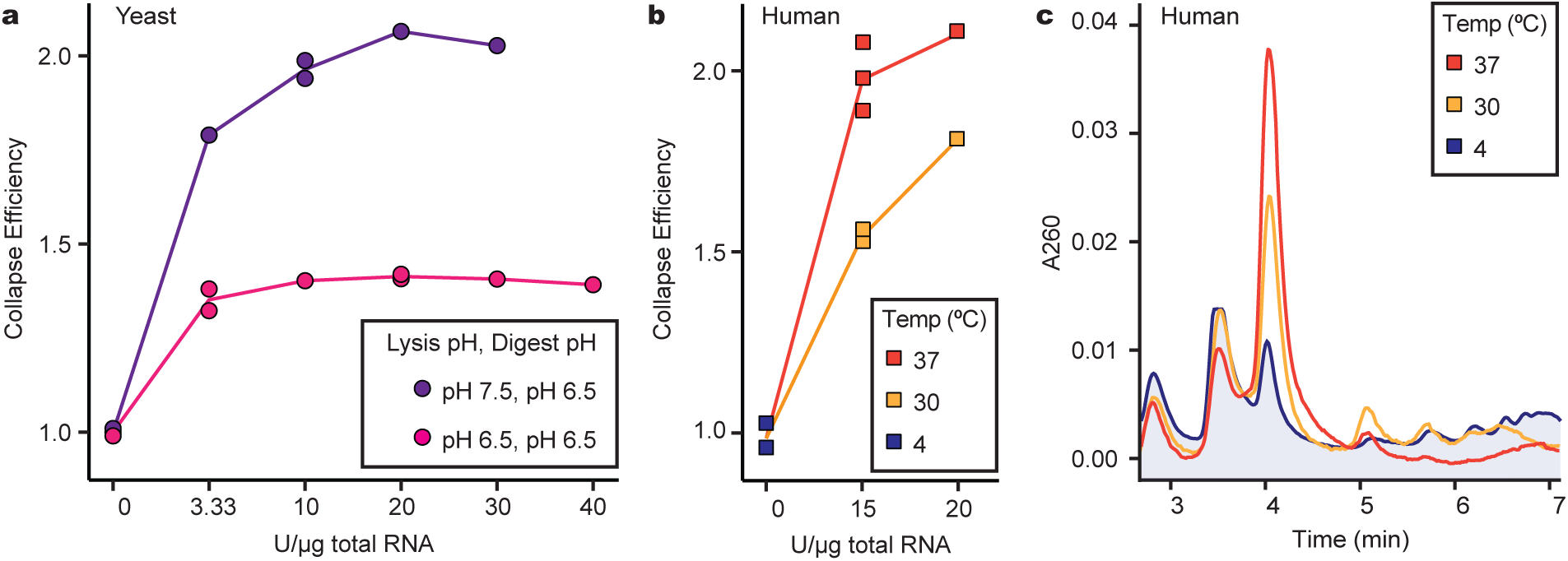
Optimization of P1 nuclease digestion and cell lysis conditions. **a**, Polysome collapse efficiency analysis comparing lysis with either pH 6.5 or pH 7.5 polysome lysis buffer and addition of a range of P1 nuclease U/µg. All assays were carried out with lysate measuring 30 µg of total RNA *quantum satis* to 200 µL with either pH 6.5 or pH 7.5 polysome buffer. At the point of nuclease digestion, pH 7.5 lysate was adjusted to ∼pH 6.5 with 14 µL of 300 mM Bis-Tris (pH 6.0), and pH 6.5 lysate was supplemented with an additional 14 µL of pH 6.5 polysome buffer. Collapse efficiency was calculated as the ratio of the integrated monosome peak absorbance relative to the integrated polysome region absorbance, normalized by the undigested control (n=1 for each condition, except n=2 for pH 6.5 with 3.33 U/µg or 20 U/µg and pH 7.5 with 0 U/µg or 10 U/µg). **b**, Comparison of polysome collapse efficiency by P1 nuclease digestion at either 30 °C or 37 °C using Calu-3 human cell lysate. Briefly, lysate measuring 15 µg of total RNA was *quantum satis* to 200 µL in pH 7.5 polysome buffer and pH adjusted with 14 µL of 300 mM Bis-Tris (pH 6.0) before supplemented with P1 nuclease and digestion at 30 °C or 37 °C. Undigested control incubated at 4 °C for an hour without nuclease (n=2 for 4 °C no-nuclease controls, n=2 or 3 for 15 U/µg as shown, and n=1 for 20 U/µg). **c**, Representative polysome profile from P1 nuclease digestion of Calu-3 human cell lysate at 30 °C or 37 °C with nuclease at 15 U/µg total RNA. Undigested control incubated at 4 °C for an hour without nuclease.

**Extended Data Figure 3:**
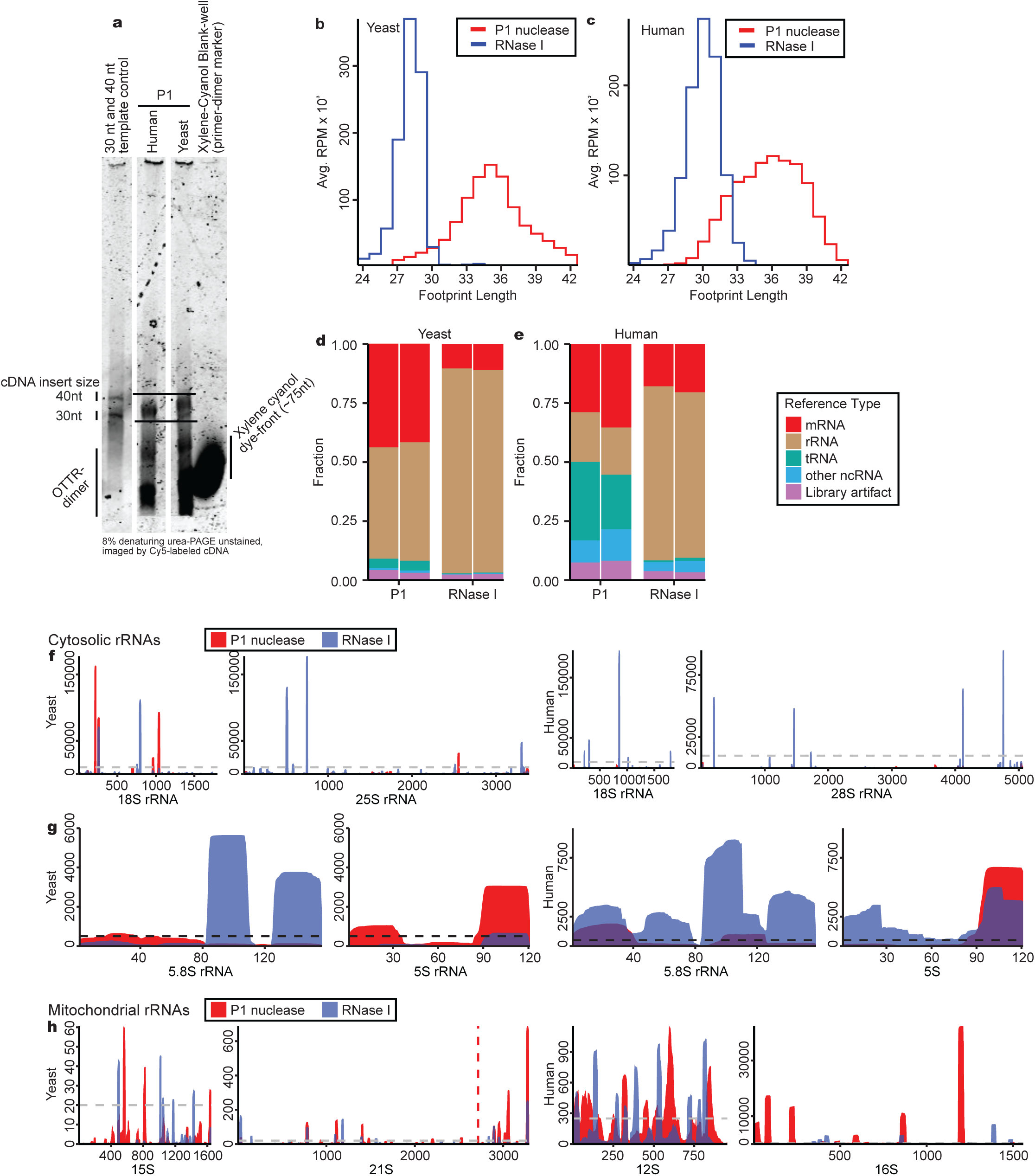
OTTR library production and ribosome profiles. **a,** Size-selection of P1 nuclease RPF cDNA from OTTR by direct imaging of Cy5, the dye covalently linked to the 5′ end of the +1dY DNA/RNA adapter duplex primer (see Fig. 1a). The 30 nt and 40 nt RNA oligonucleotides used for RPF size selection (not shown) were also used in OTTR reactions parallel to those using input RPFs, to generate cDNA size-selection markers. Bromophenol blue formamide loading dye was used to resuspend OTTR cDNA for size selection to avoid xylene cyanol interference during Cy5 imagining. A 0.6X TBE 8% urea-PAGE was chosen for cDNA size selection since xylene cyanol and the no-insert OTTR adapter-dimer cDNA (∼75 nt) co-migrate. For these reasons, xylene cyanol was included only in the peripheral lanes. Horizontal black lines indicate the boundaries for cDNA gel slice excision to remove adaptor-dimer from desired cDNA library. All lanes are from the same gel. P1 RPF were either from human 293T or S288C yeast cell lysate. A 30 nt and 40 nt template control OTTR reaction was used to synthesize OTTR cDNA to enable cDNA size selection equivalent to RNA size selection. **b**, Read length distribution of yeast RNase I (blue) and P1 nuclease (red) RPFs from the CDS, excluding those RPFs that are aligned to the first 15 and last 10 codons, as in Extended Data Fig. 1b. Counts were represented in RPM and averaged across replicates **c**, Read length distribution of human RNase I (blue) and P1 nuclease (red) RPFs from the CDS, excluding those RPFs that are aligned to the first 15 and last 10 codons, as in Extended Data Fig. 1b. Counts were represented in RPM and averaged across replicates **d**, Fraction of RNase I RPF cDNA library sequencing reads mapped to each transcript class for yeast libraries generated by P1 nuclease or RNase I digestion. **e**, Fraction of RNase I RPF cDNA library sequencing reads mapped to each transcript class for human libraries generated by P1 nuclease or RNase I digestion. **f**, Average per-base read coverage of cytosolic 18S and 25S or 28S rRNA from yeast (left) or human (right) ribosome profiles with P1 nuclease (red) or RNase I (blue). Coverage was represented in reads per million total reads, including reads mapping to rRNA, tRNA, ncRNA, mRNA, and other genomic loci) to emphasize relative proportion from the entire library. **g**, As in (**f**) but for 5.8S and 5S rRNA coverage. **h**, As in (**f**) for mitochondrial rRNA coverage.

**Extended Data Figure 4:**
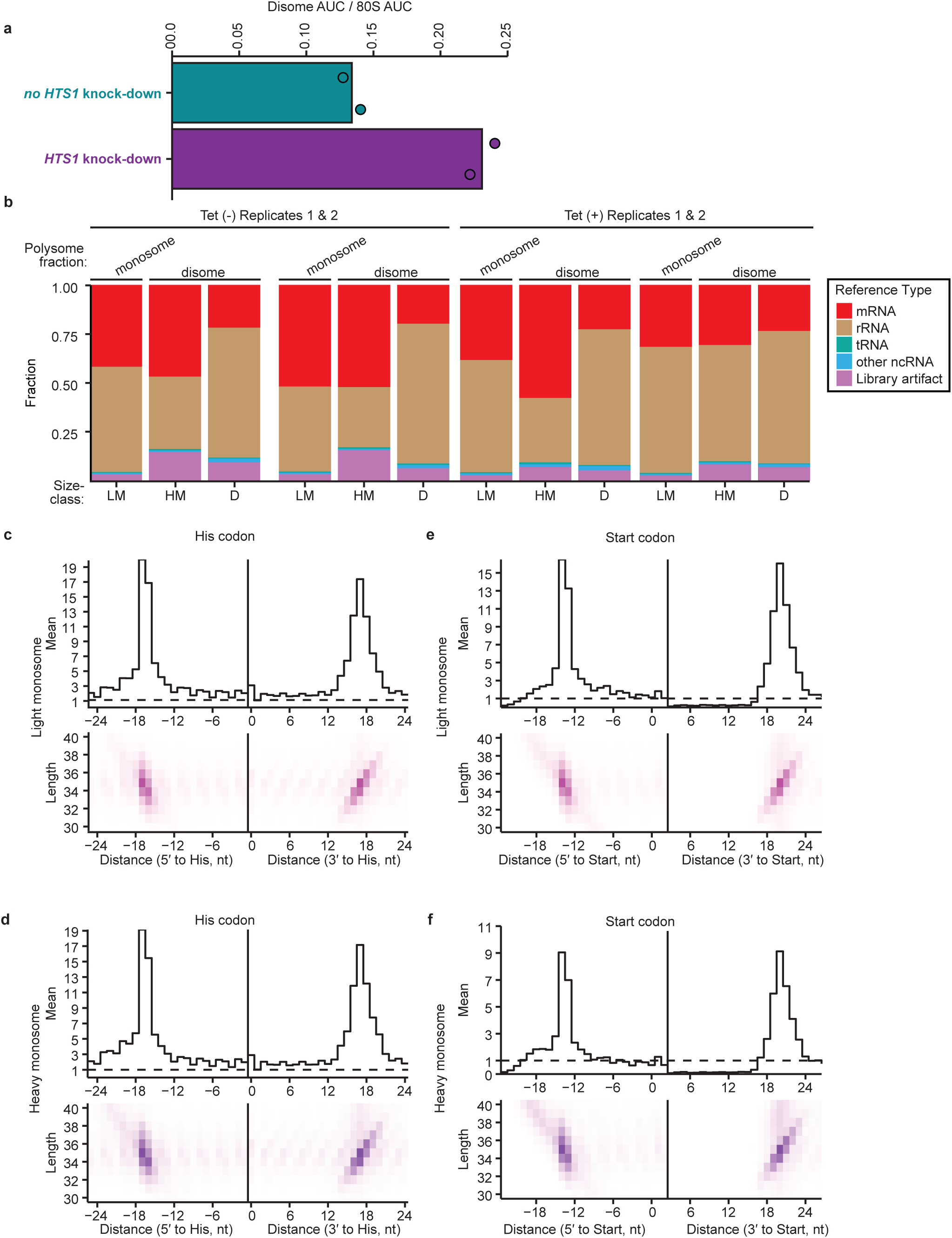
Abundance and properties of monosome, sub-disome, and true disome RPFs from yeast cells with or without histidine starvation. **a**, Ratio of disome to monosome peak area from polysome profiles after P1 nuclease digestion of yeast lysates from cells with or without *HTS1* knockdown. **b**, Fraction of sequencing reads mapped to each transcript class for light monosome, heavy monosome, and disome profiling from P1 nuclease digestion and OTTR library cDNA synthesis. **c**, Average profile of footprints around isolated histidine codons after *HTS1* depletion, as in Fig. 4g but for light monosome RPFs. **d**, Average profile of footprints around isolated histidine codons after *HTS1* depletion, as in Fig. 4g but for heavy monosome RPFs. **e**, Average profile of footprints for light monosome RPFs at start codons after *HTS1* depletion **f**, Average profile of footprints for heavy monosome RPFs at start codons after *HTS1* depletion

**Extended Data Figure 5:**
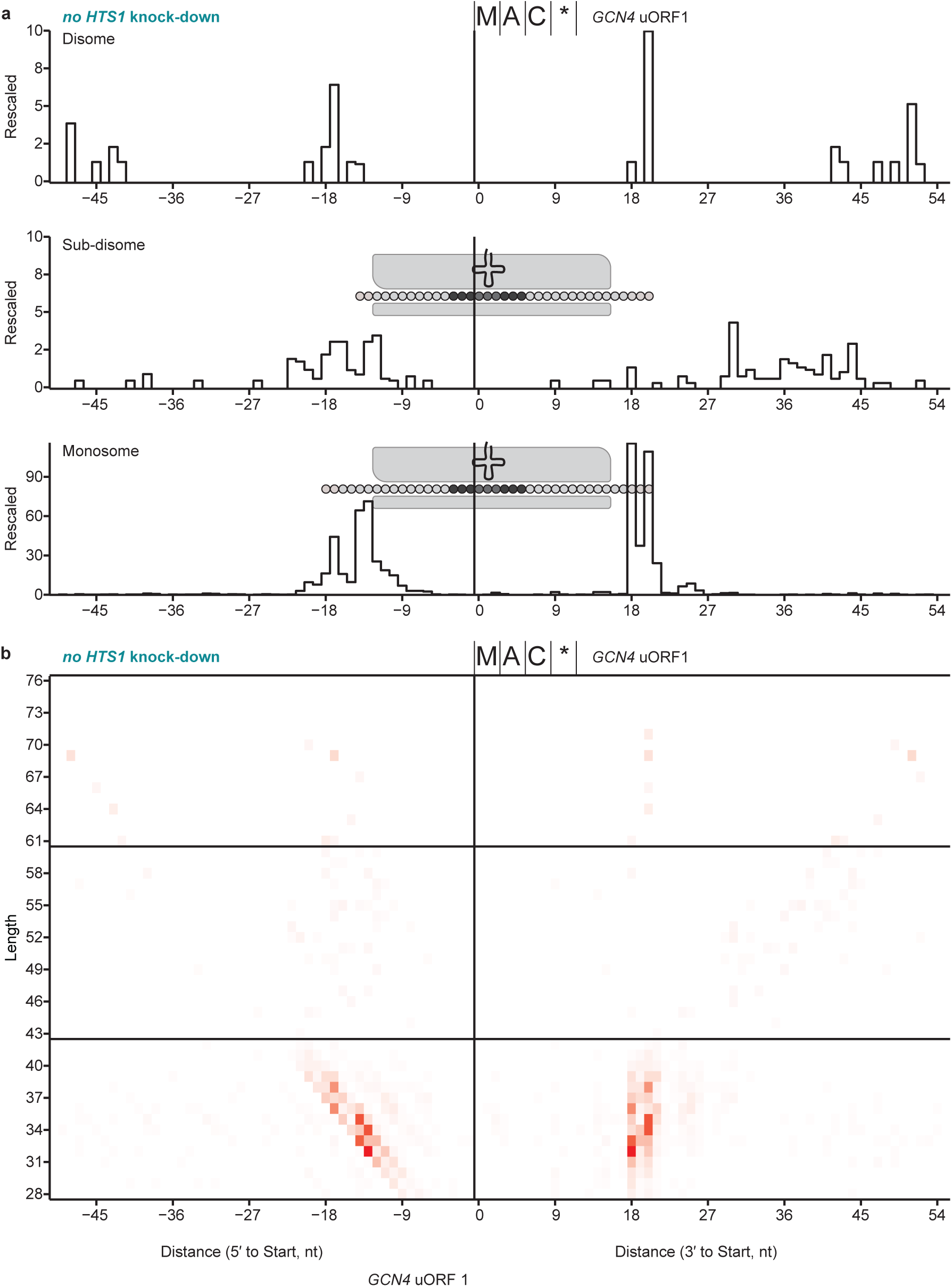
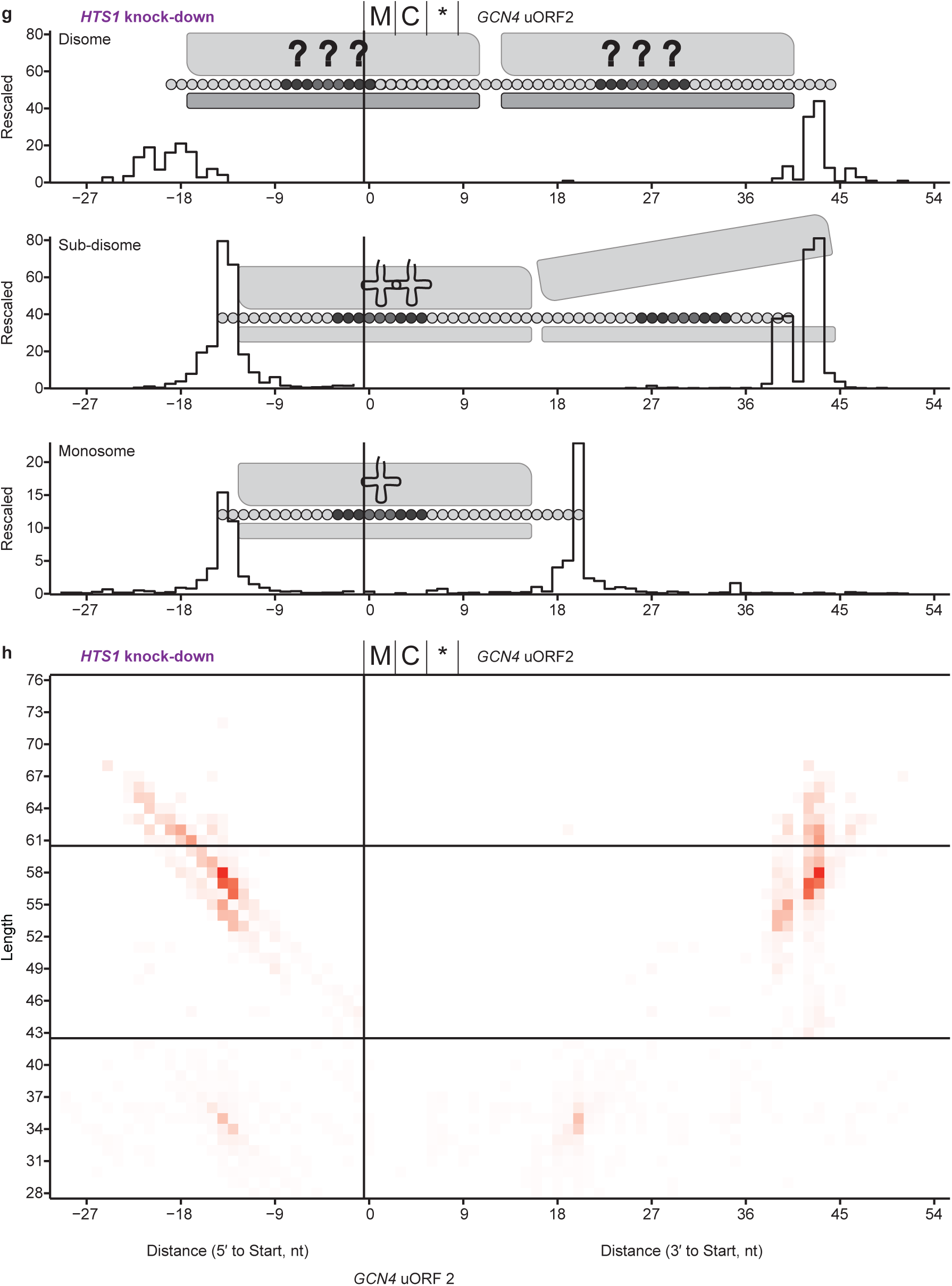
Complete *GCN4* uORF1 and uORF2 profiles for monosome, sub-disome, and true disome footprints with and without *HTS1* knock-down. **a**, Extended representation from Figure 5c. Rescaled counts of 5′ and 3′ ends of aligned true disome (top), sub-disome (middle), and monosome (bottom) footprints for *GCN4* uORF1 without *HTS1* knock-down. Results from replicates were summed together. **b**, Footprint length 5′ and 3′ profile of (**a**). **c**, Rescaled counts of 5′ and 3′ ends of aligned true disome (top), sub-disome (middle), and monosome (bottom) footprints for *GCN4* uORF1 after *HTS1* knock-down. Results from replicates were summed together. **d**, Footprint length 5′ and 3′ profile of (**c**). **e**, Extended representation from Figure 5d. Rescaled counts of 5′ and 3′ ends of aligned true disome (top), sub-disome (middle), and monosome (bottom) footprints for *GCN4* uORF2 without *HTS1* knock-down. Results from replicates were summed together. **f**, Footprint length 5′ and 3′ profile of (**e**). **g,** Rescaled counts of 5′ and 3′ ends of aligned true disome (top), sub-disome (middle), and monosome (bottom) footprints for *GCN4* uORF2 after *HTS1* knock-down. Results from replicates were summed together. **h**, Footprint length 5′ and 3′ profile of (**g**).

**Extended Data Figure 6:**
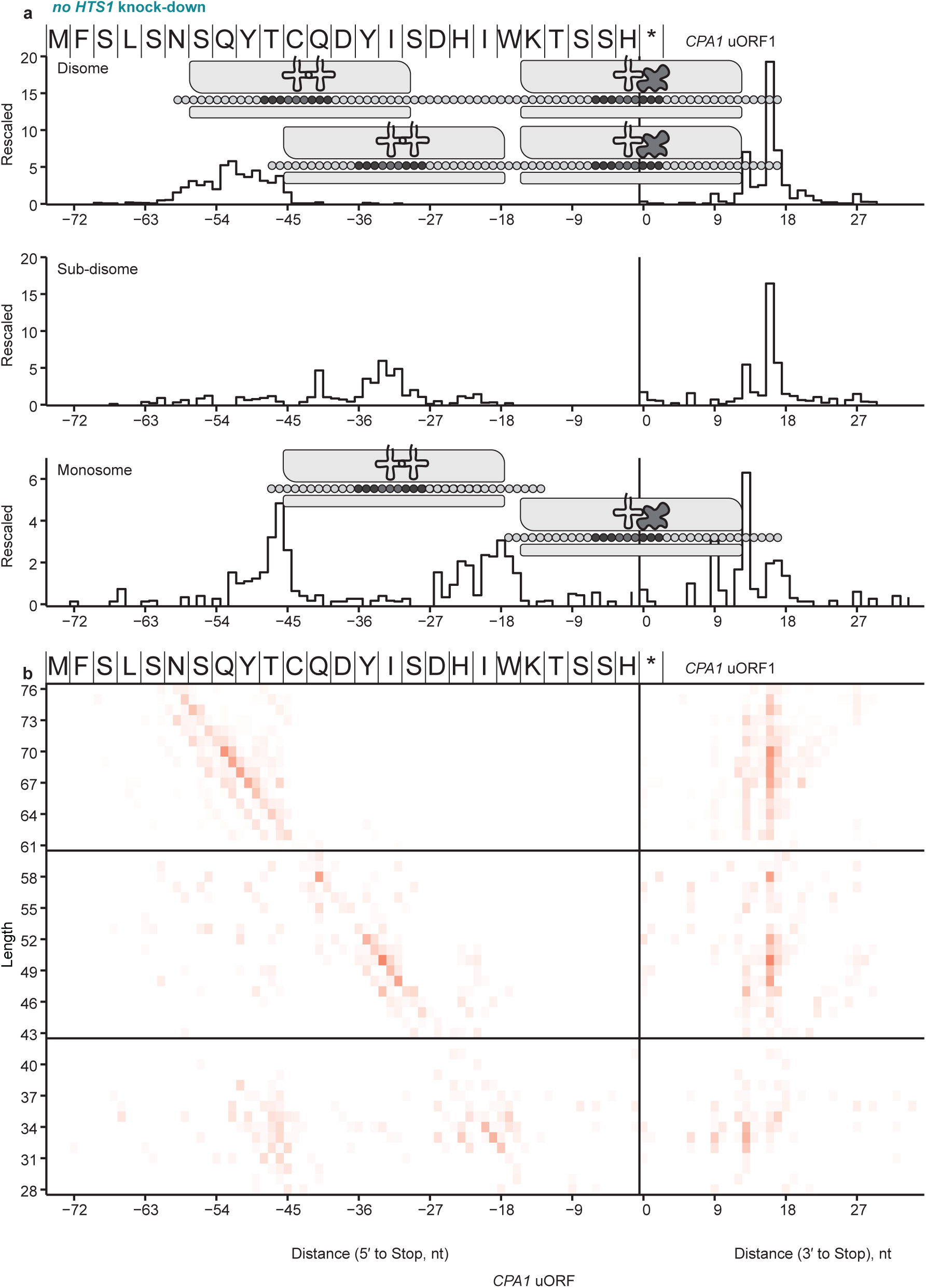
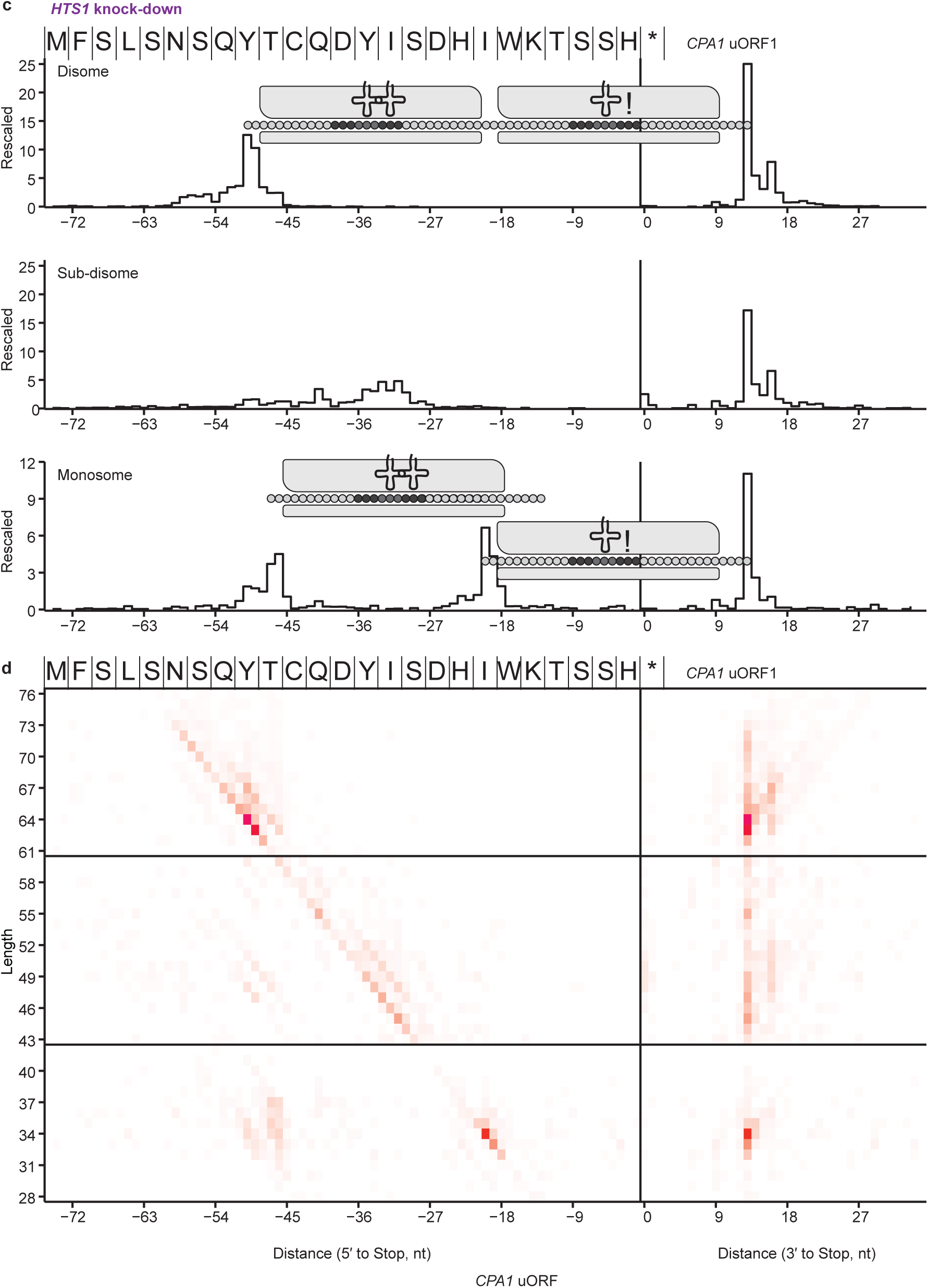
Complete *CPA1* uORF profiles for monosome, sub-disome, and true disome footprints with and without *HTS1* knock-down. **a**, Extended representation from Figure 6a. Rescaled counts of 5′ and 3′ ends of aligned true disome (top), sub-disome (middle), and monosome (bottom) footprints for *CPA1* uORF without *HTS1* knock-down. Results from replicates were summed together. **b**, Extended representation from Figure 6b. Rescaled counts of 5′ and 3′ ends of aligned true disome (top), sub-disome (middle), and monosome (bottom) footprints for *CPA1* uORF after *HTS1* knock-down. Results from replicates were summed together.

**Extended Data Figure 7:**
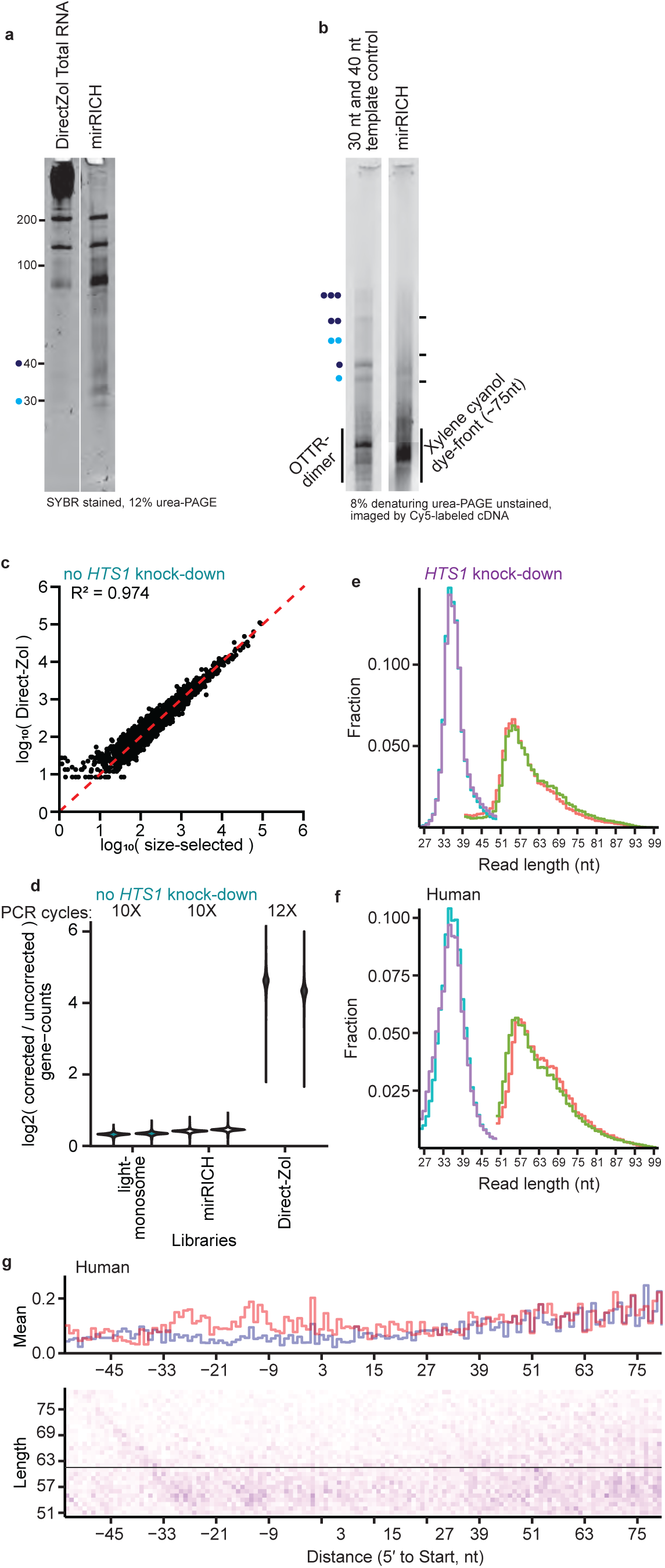
Additional comparisons of libraries made from mirRICH, total RNA, or gel-base size-selected RPFs. **a**, Denaturing urea-PAGE analysis of yeast RNA fragments purified from either Direct-Zol Total RNA extraction or mirRICH small RNA enrichment. Roughly 5% of the Direct-Zol extract and 25% of the mirRICH extract was analyzed in this gel. The migration of the 30 nt (light blue) and 40 nt (dark blue) size-selection RNA oligos are demarcated (left). The gel was a 12% urea-PAGE with 0.6X TBE, and SYBR gold was used for staining. Both lanes are from the same gel. **b**, Denaturing urea-PAGE analysis of cDNA produced from OTTR using 40 ng of mirRICH small RNA. Parallel positive control OTTR libraries were synthesized from either the 30 nt or 40 nt RNA size-selection RNA oligos to aid in cDNA size-selection. The single light-blue and dark-blue dots (left) demarcate the cDNA with inserts derived from the 30 nt and 40 nt RNA oligos, respectively. cDNA concatemers in OTTR can form if excess template is included in the reaction, such as the 60 nt and 80 nt sized-inserts demarcated by two light blue and two dark blue dots, respectively. The approximate area for monsome cDNA size selection was demarcated by the bottom two black lines (right), and the disome cDNA was size selected from the top two black lines. The gel was an 8% urea-PAGE with 0.6X TBE, and cDNA was directly imaged by Cy5. Both lanes are from the same gel. **c**, Gene-level DESeq2 estimates for CDS occupancy (as in Figure 1b), but here for mirRICH or Direct-Zol P1 RPF from the yeast lysate without *HTS1* knock-down. Both libraries relied on cDNA size-selection rather than RNA size-selection. The counts were deduplicated before analysis, and a gene’s CDS needed at least one alignment from one replicate in order to be retained for analysis. **d**, Distribution of the log_2_ ratio of PCR deduplicated counts (corrected) versus uncorrected counts for each CDS from the libraries generated from the no *HTS1* knock-down lysate. The number of necessary PCR cycles used for library multiplexing is defined above. Two technical replicates per library condition were analyzed. **e**, Read length distribution represented as a fraction of mRNA mapping reads for the mirRICH monosome and disome cDNA size-selected libraries from the *HTS1* knock-down lysate. Two technical replicates per library condition were analyzed. Monosome replicates are in light blue and purple; disome replicates are in red and green. **f**, Read length distribution represented as a fraction of mRNA mapping reads for the mirRICH monosome and disome cDNA size-selected libraries from the 293T lysate. Two technical replicates per library condition were analyzed. Monosome replicates are in light blue and purple; disome replicates are in red and green. **g**, 5′ aligned ends and footprint length profile at initiating codons for P1 sub-disome and true disome RPFs captured by mirRICH from human 293T cell lysates. The summed 5′ end profile is depicted at top and the contributions from each footprint length at each 5′ end are shown at bottom. Sub-disome and true disome sized reads were analyzed separately.

